# Alterations in protein *N*-glycosylation confer vanadate resistance in *Ogataea polymorpha* mutants defective in phosphomannosylation

**DOI:** 10.1101/2025.11.05.686784

**Authors:** Maria Pakhomova, Azamat Karginov, Maria Kulakova, Polina Vladimirova, Olga Mitkevich, Michael Agaphonov

**Affiliations:** The Federal Research Center “Fundamentals of Biotechnology” of the Russian Academy of Sciences, 119071 Moscow, Russia

**Keywords:** protein secretion, protein glycosylation, phosphomannosylation, vanadate resistance, *Ogataea*, yeast

## Abstract

Different yeast species including *Ogataea polymorpha* are often used as hosts for recombinant protein production. One of the most important factors limiting such applications is yeast-specific modifications of glycoside chains attached to secretory proteins. This problem can potentially be solved by the identification and inactivation of genes responsible for these modifications. Previously we demonstrated that the exceptional resistance of *O. polymorpha* to vanadate depends on the *ABV1* gene responsible for the mannosylphosphorylation of protein glycoside chain in the Golgi apparatus. Here we show that mutations altering protein glycosylation in the secretory pathway can be selected in the *abv1-Δ* mutant by screening for vanadate resistance. For one such mutant, we identified the responsible gene, which encodes a putative α-1,2-mannosyltransferase. To ensure the absence of phosphomannosylation, both *O. polymorpha* genes, *ABV1* and *MNN4*, which encode mannosylphosphate transferase homologs, were inactivated. Some vanadate resistant mutants generated in this strain showed defects in *N*-glycosylation of a recombinant glycoprotein. This demonstrates that the effects of *N*-glycosylation on vanadate resistance in *O. polymorpha* are not mediated by phosphomannosylation per se and that identification of certain genes responsible for *N*-glycosylation in this yeast can be performed via selection of vanadate resistant clones.

## Introduction

Proteins that enter the eukaryotic secretory pathway may undergo different posttranslational modifications, one of which is *N*-glycosylation. It starts in the endoplasmic reticulum (ER) with the attachment of Glc_3_Man_9_GlcNAc_2_ oligosaccharide to specific asparagine residues in the -N-X-S/T- motif, where X is any amino acid except proline. This oligosaccharide is processed to the Man_8_GlcNAc_2_ structure before the protein enters the Golgi apparatus, where the glycoside receives additional modifications that can be essentially different in different organisms. In *Saccharomyces cerevisiae*, this Man_8_GlcNAc_2_ core glycoside can be modified by the attachment of a branched chain of mannose residues, a process initiated by the Och1 α-1,6-mannosyltransferase. However, the core glycoside of some *S. cerevisiae* proteins, e.g. carboxypeptidase Y, receive only few additional mannose residues (for a review see (Munro, 2001)). These modifications are frequently undesirable when yeast cells are used as a host for production of recombinant glycoproteins. To solve this problem it is necessary to inactivate the host genes encoding specific mannosyltransferases and introduce some heterologous genes, which encode enzymes catalyzing the desired modifications (for review see (Madhavan et al., 2021; Li et al., 2022)).

Methylotrophic yeasts such as *Komagataella phaffii* (syn. *Pichia pastoris*) and two species of the *Ogataea* genus, *O. polymorpha* and *O. parapolymorpha*, which were formerly classified as *Hansenula polymorpha*, are frequently used as hosts for highly efficient recombinant protein production (Manfrão-Netto et al., 2019; Barone et al., 2023). Proteins secreted by these yeasts are usually not hyperglycosylated and their *N*-linked glycoside chains possess 8-14 mannose residues (Bretthauer and Castellino, 1999; Kim et al., 2004). While the *K. phaffii* homolog of *S. cerevisiae* Och1 was shown to be responsible for the initiation of mannosylation of the core glycoside in the Golgi apparatus (Choi et al., 2003), the closest homolog of this protein in *O. parapolymorpha* may not play a key role in this process. Functional analysis of other genes encoding homologs of α-1,6-mannosyltransferase in the latter yeast species identified the *OCR1* gene whose inactivation prevented the attachment of α-1,6-mannose to the core glycosides of a recombinant protein in the Golgi apparatus (Kim et al., 2006).

In yeast, in addition to mannose residues, *N*- and *O*-linked glycoside chains are also modified by attachment of phosphomannose (Jigami and Odani, 1999). This modification of cell-wall mannoproteins creates negative charge on the cell surface that can be revealed by the ability of cells to bind Alcian blue dye. In *S. cerevisiae*, phosphomannose attachment depends on the protein encoded by the *MNN4* gene (Jigami and Odani, 1999), which belongs to the fukutin protein family and presumably catalyzes mannosylphosphate transfer from the GDP-mannose donor (Aravind and Koonin, 1999). There is an *MNN4* paralog in the *S. cerevisiae* genome, designated *MNN14*, however the inactivation of *MNN4* alone is sufficient to abolish the Alcian blue staining (Jigami and Odani, 1999).

It is well known that mutations conferring resistance to vanadate in *S. cerevisiae* frequently affect protein *N*-glycosylation (Kanik-Ennulat et al., 1995), however the mechanism of this effect is not clear. Vanadate is a potent inhibitor of a large variety of enzymes due to its similarity to phosphate (Stankiewicz et al., 1995). These multiple targets can be protected by restriction of vanadate uptake from the environment. Vanadate resistant mutants isolated in *Candida albicans* (Mahanty et al., 1991) and Neurospora crassa (Bowman, 1983; Bowman et al., 1983) were defective in phosphate transport. The yeast phosphate transport system includes low- and high-affinity transporters. *S. cerevisiae* possesses two highly homologous low-affinity transporters in the plasma membrane, namely Pho87 and Pho90, and two high-affinity transporters Pho84 and Pho89, which show no similarity. Additionally, there is the vacuolar membrane low-affinity phosphate transporter Pho91. Despite high similarity, Pho87 and Pho90 are not equally functional, since the former is more efficient as a phosphate sensor, while the latter is more efficient as a transporter. The high-affinity transporter Pho84 was also sown to have an external phosphate sensing function (Giots et al., 2003). Methylotrophic yeast *Ogataea polymorpha* and *O. parapolymorpha* possess only one gene for the plasma membrane low-affinity phosphate transporter also designated as Pho87. Its inactivation in *O. parapolymorpha* increases resistance to vanadate (Karginov et al., 2018), while inactivation of the Pho91 homolog does not affect this phenotype (Farofonova et al., 2024). The *O. parapolymorpha* low-affinity phosphate transporters are somehow related to the regulation of methanol oxidase activity since inactivation of Pho91 leads to its complete reduction (Farofonova et al., 2023) while additional inactivation of Pho87 restores it to the wild-type level (Farofonova et al., 2024).

In contrast to *O. parapolymorpha*, whose vanadate resistance is similar to that of *S. cerevisiae* (Kim et al., 2013), its closest relative *O. polymorpha* is highly resistant to this compound (Mannazzu et al., 1997). Screening for vanadate hypersensitive mutants in this yeast identified the gene designated *ABV1* (Karginov et al., 2018), which encodes a protein homologous to the *S. cerevisiae* putative mannosylphosphate transferase Mnn4. Its inactivation in both *O. polymorpha* and *O. parapolymorpha* led to a drastic decrease in the Alcian blue staining indicating defect in mannosylphosphorylation of oligosaccharide chains of cell-wall glycoproteins (Karginov et al., 2018). These yeasts possess one more gene encoding protein, which is even more homologous to *S. cerevisiae* Mnn4, but is probably not generally involved in mannosylphosphorylation of oligosaccharides of cell wall proteins, since inactivation of *ABV1* alone was sufficient to virtually abolish Alcian blue staining. This gene is annotated as *MNN4* in the deposited in NCBI GenBank *O. parapolymorpha* genome sequence (Ravin et al., 2013) due to its high similarity to the *S. cerevisiae MNN4*.

The role of *O. polymorpha* Abv1 in vanadate resistance most probably is related to the regulation of expression of the gene for the high-affinity transporter Pho84, since deletion of the *ABV1* gene activates the *PHO84* promoter while increase in *ABV1* dosage represses it (Karginov et al., 2018). This may explain why *S. cerevisiae* mutants defective in glycosylation frequently show increased vanadate resistance, if the same mechanism is relevant in this yeast. At the same time, it remains unclear whether mannosylphosphorylation per se is crucial for vanadate resistance, or other alterations of *N*-linked glycoside chains can also affect this trait even in the absence of Abv1 function. The aim of this study was to explore these possibilities as well as to identify *O. polymorpha* genes affecting vanadate resistance and altering the protein glycosylation in the secretory pathway.

## Materials and Methods

### Yeast strains, transformation and culture conditions

The strains used in this study are listed in Table 1. The *O. polymorpha* strain M257 (*leu2 zeo*^*R*^ *P*_*MOX*_*-GOX*) carrying an expression cassette of *Aspergillus niger* glucoseoxidase (GOX) and 1b27 (*leu2 ade2 ura3::ADE2*) were used as a wild-type strain to construct mutants defective in protein glycosylation. The 1B27 was described previously (Fokina et al., 2012). The M257 strain was obtained by transformation of the A16 strain (Veale et al., 1992) with the linearized pZAM518 plasmid, which carries the GOX expression cassette under the control of the *MOX* promoter and the zeocin resistance selectable marker. Yeast strains were transformed as described previously (Karginov et al., 2021). Synthetic complete medium with glucose as a carbon source (0,67% Yeast Nitrogen Base, 2% glycose, 2% agar) was used to select yeast transformants. YPD (1% Yeast Extract, 2% Peptone, 2% Glucose) and YPM (1% Yeast Extract, 2% Peptone, 1% Methanol) media were used to cultivated yeast strains. To induce GOX expression, yeast cells were grown in Y3PM (1% Yeast Extract, 3% Peptone, 1% Methanol) supplemented with 150 mM NaCl at 30°C for 50-60 h.

**Table 1.** Yeast strains used in this study.

| Strain | Genotype |
| --- | --- |
| M257 | *leu2 zeo<sup>R</sup> P<sub>MOX</sub>-GOX* |
| M257-620M1 | *leu2 zeo<sup>R</sup> P<sub>MOX</sub>-GOX abv1::loxP* |
| M257-620-759M1 | *leu2 zeo<sup>R</sup> P<sub>MOX</sub>-GOX abv1::loxP abv2::loxP* |
| M257-MP6 | *leu2 zeo<sup>R</sup> P<sub>MOX</sub>-GOX och1::LEU2* |
| M257-1055 | *leu2 zeo<sup>R</sup> P<sub>MOX</sub>-GOX ocr1::loxP* |
| M257-MP9 | *leu2 zeo<sup>R</sup> P<sub>MOX</sub>-GOX mnn4-Δ* |
| M257-620-MP9 | *leu2 zeo<sup>R</sup> P<sub>MOX</sub>-GOX abv1::loxP mnn4-Δ* |
| M257-620-MP6 | *leu2 zeo<sup>R</sup> P<sub>MOX</sub>-GOX abv1::loxP och1::LEU2* |
| M257-620-MP9-MP6 | *leu2 zeo<sup>R</sup> P<sub>MOX</sub>-GOX abv1::loxP mnn4-Δ och1::LEU2* |
| M257-759M1 | *leu2 zeo<sup>R</sup> P<sub>MOX</sub>-GOX abv2::loxP* |
| 1B27 | *leu2 ade2 ura3::ADE2* |
| 1B27-620M1 | *leu2 ade2 ura3::ADE2 abv1::loxP* |

### Plasmids, oligonucleotides and gene disruption cassettes

Oligonucleotides used in this work are listed in Table S1. The pZAM518 plasmid was based on the pDLMOX-GOD-H GOX expression vector (Kim et al., 2004), whose 2405 bp DraI-BamHI fragment bearing the ampicillin resistance and *LEU2* selectable markers was replaced with the 923 bp BamHI-EcoRV fragment of the pGAPZα-A plasmid (Invitrogen) carrying the zeocin resistance marker. Construction the pAM620 plasmid used to disrupt the *ABV1* gene with *cre/loxP*-self excisable vector was described previously (Karginov et al., 2018). To obtain recombination arms for the *ABV2* disruption cassette, *O. polymorpha* genomic DNA was digested with EcoRV, self-ligated and used as a template for PCR with primers ABV2U and ABV2L. The obtained fragment was inserted between the EcoRV and the SalI sites of the pAM619 vector (Agaphonov and Alexandrov, 2014). The resulting plasmid designated pAM759 was linearized with EcoRV and used for yeast transformation to disrupt the *ABV2* gene. To recycle the *leu2* auxotrophic marker, the disruptants obtained with plasmids pAM620 and pAM759 were grown overnight on liquid YPM and spread onto YPD plates to obtain separate colonies, which were then tested for the loss of leucine prototrophy.

Since there is the *HIS2* gene adjacent to *MNN4* in the *O. polymorpha* genome it could be used to select transformants with plasmid integrated into this locus. Thus, we have constructed two plasmids for the *MNN4* disruption, one of which, pMP9V, was designed to replace *MNN4* and portion of *HIS2* with the *LEU2* selectable marker, and another one pCMP7 designed to replace integrated sequence of pMP9V bearing *LEU2* and to restore *HIS2*. To construct the pCMP7 plasmid, one recombination arm was obtained by PCR with primers OpoMNN4AU1 and OpoMNN4L1 and digested with HindIII, while the other was obtained by PCR with primers OpoMNN4AL1 and OpoMNN4U1 and digested with BamHI and HindIII. The obtained fragments were ligated with the BamHI-HincII-digested pBCKS+ vector. To construct pMP9V plasmid, the 2963 bp ScaI Eco72I fragment of pCMP7 was inserted between the EcoRV and the BglII sites of the pAM773 vector (Agaphonov, 2017).

The *OCH1* disruption cassette was constructed as follows. The DNA fragment representing the *O. polymorpha* genomic locus carrying *OCH1* was obtained by PCR with primers OpoOCH1U1 and OpoOCH1L1, digested with HindIII and ligated with HindIII-Ecl136II-digested pUC18 vector. The NaeI-BglII 241 bp fragment within the *OCH1* ORF in the resulting plasmid was replaced with the BamHI-EcoRV 1254 bp fragment of the pCLHX plasmid (Sohn et al., 1996) carrying the *LEU2* selectable marker. The resulting plasmid designated, which was designated pMP6, was digested with HindIII and NcoI to excise the disruption cassette used to disrupt *OCH1* in the *O. polymorpha* genome.

Sequences of the *O. parapolymorpha OCR1* locus were used as recombination arms for the *OCR1* disruption cassette. To construct it, the DNA fragment possessing this gene was obtained by PCR with primers OpaOCR1AL and OpaOCR1AU. The BamHI-PstI 1382 bp and XhoI-PstI 1316 bp were ligated with the BamH-XhoI-digested pBCKS+ plasmid to obtain pAM1042. The recombination arms were excised from pAM1042 by BssHII-BamHI (971bp fragment) and BssHII-PvuII (1346bp fragment) and ligated with the EcoRV-BglII-cleaved pAM773 vector, which is capable of self-excision by cre/loxP recombination (Agaphonov, 2017). The resulting plasmid designated pAM1055 was cleaved with XhoI and BglII prior to yeast transformation.

### UV mutagenesis

To obtain vanadate resistant mutants, yeast cells from logarithmic YPD culture were collected by centrifugation re-suspended in sterile water to OD_600_ ∼0.5 and exposed to UV for 10”. The cell suspension was mixed with an equal volume of YPD and incubated in 37°C shaker incubator for 1h. Cells were collected by centrifugation, re-suspended in YPD containing 17% glycerol dispensed by 0.5 ml in 1.5 ml Eppendorf tubes and stored at -70°C. The frozen aliquots were thawed at room temperature and spread onto YPD plates containing 3.5 or 4 mM sodium orthovanadate.

### Electrophoresis, immunoblotting and antibodies

Proteins were separated by SDS-PAGE and transferred to PVDF membrane according to standard protocols (Laemmli, 1970; Towbin et al., 1979). Commercial antibody against *Aspergillus niger* glucose oxidase (Accurate Chemical & Scientific, Westbury, NY) was kindly provided by prof. H.-A. Kang (Korea). Rabbit antiserum against *E. coli-*expressed *O. polymorpha* chitinase was used to detect this protein in culture supernatants. *N*-glycoside chains were removed from GOX by treatment with recombinant PNGase F (Boyko et al., 2023), which was kindly provided by Dr. N. Sluchanko (Moscow, Russia).

### Alcian blue staining

Cell staining with Alcian blue was performed as described previously (Ballou, 1990) with minor modifications as follows. Cells from overnight cultures were precipitated by centrifugation at 2500 g for 3 min, washed with 10 mM HCl and re-suspended in 0.1% Alcian blue solution in 10 mM HCl. Unbound dye was removed by washing cells with 10 mM HCl. Cells were transferred into a flat-bottom 96-well plate precipitated by centrifugation (2000 g, 3 min) and scanned.

## Results

### Mutations affecting *N*-linked glycosylation can increase vanadate resistance in *O. polymorpha abv1-Δ* mutant

As it was mentioned in the Introduction, inactivation of the *O. polymorpha ABV1* gene, which encodes mannosylphosphate transferase, decreases vanadate resistance (Karginov et al., 2018). To explore whether other mutations can increase vanadate resistance when this gene is deleted, the 1B27-620M1 strain possessing a deletion *abv1* allele was UV-mutagenized and vanadate resistant clones were selected on YPD containing 4 mM sodium orthovanadate. Unlike the original strain, one of the obtained vanadate resistant clones, AV35, was unable to grow at 45°C and in presence of 0.005% SDS in culture medium. As it was mentioned in the introduction, it was known that in *S. cerevisiae* some mutations affecting *N*-glycosylation also increase vanadate resistance (Kanik-Ennulat et al., 1995), while defects of glycosylation in the secretory pathway in *Ogataea* yeasts were shown to increase detergent sensitivity (Agaphonov et al., 2001, 2005; Kim et al., 2006). Thus, the phenotypes observed in the AV35 mutant suggested possible defects in protein *N*-glycosylation. However the strain used for mutagenesis did not possess a convenient reporter for the analysis of *N*-glycosylation. To solve this problem, we constructed an *abv1-Δ* mutant strain expressing *A. niger* extracellular glucose oxidase (GOX) as an *N*-glycosylation reporter. This strain was also UV-mutagenized and vanadate resistant clones were selected on plates containing 3,5 mM sodium orthovanadate. Some of them grew very slowly even in the absence of vanadate, which hampered further analysis. Finally, 26 mutants with acceptable growth rates were chosen to test whether they were altered in protein *N*-glycosylation. To do this electrophoretic mobility of secreted GOX was studied. As expected, GOX migrated on the SDS PAGE as a smear due to the irregular size of *N*-glycoside chains. Electrophoretic mobility of this smear in some mutants differed from that in the original *abv1-Δ* strain (Figure S1, “AV” mutants) that indicated alterations in *N*-glycosylation. To confirm that this was due to the difference in the size of *N*-linked glycosides, electrophoretic mobility of glycosylated and deglycosylated GOX was studied in some mutants (Figure 1a and b). Unlike the untreated GOX migrating as a smear, which was different in different strains, the deglycosylated GOX migrated as two very close sharp bands (probably due to different proteolytic processing) with the same apparent molecular weight in all strains, which virtually corresponded to the calculated molecular weight of the mature polypeptide (64 kDa). This demonstrated that the difference in the mobility of the untreated GOX in different strains was due to alterations in *N*-linked glycosylation. Notably, there were less GOX species with lower electrophoretic mobility in some cases, particularly, in mutants AV13 and AV21 and this protein migrated as a more compact smear that in the original *abv1-Δ* strain indicating the decrease in *N*-glycoside chain size, while a noticeable increase in *N*-glycoside chain size was observed in the AV20 mutant (Figure 1a). These results demonstrate that modifications of glycoside chains other than that catalyzed by the Abv1 protein can also affect vanadate resistance in *O. polymorpha*.

**Figure 1.**
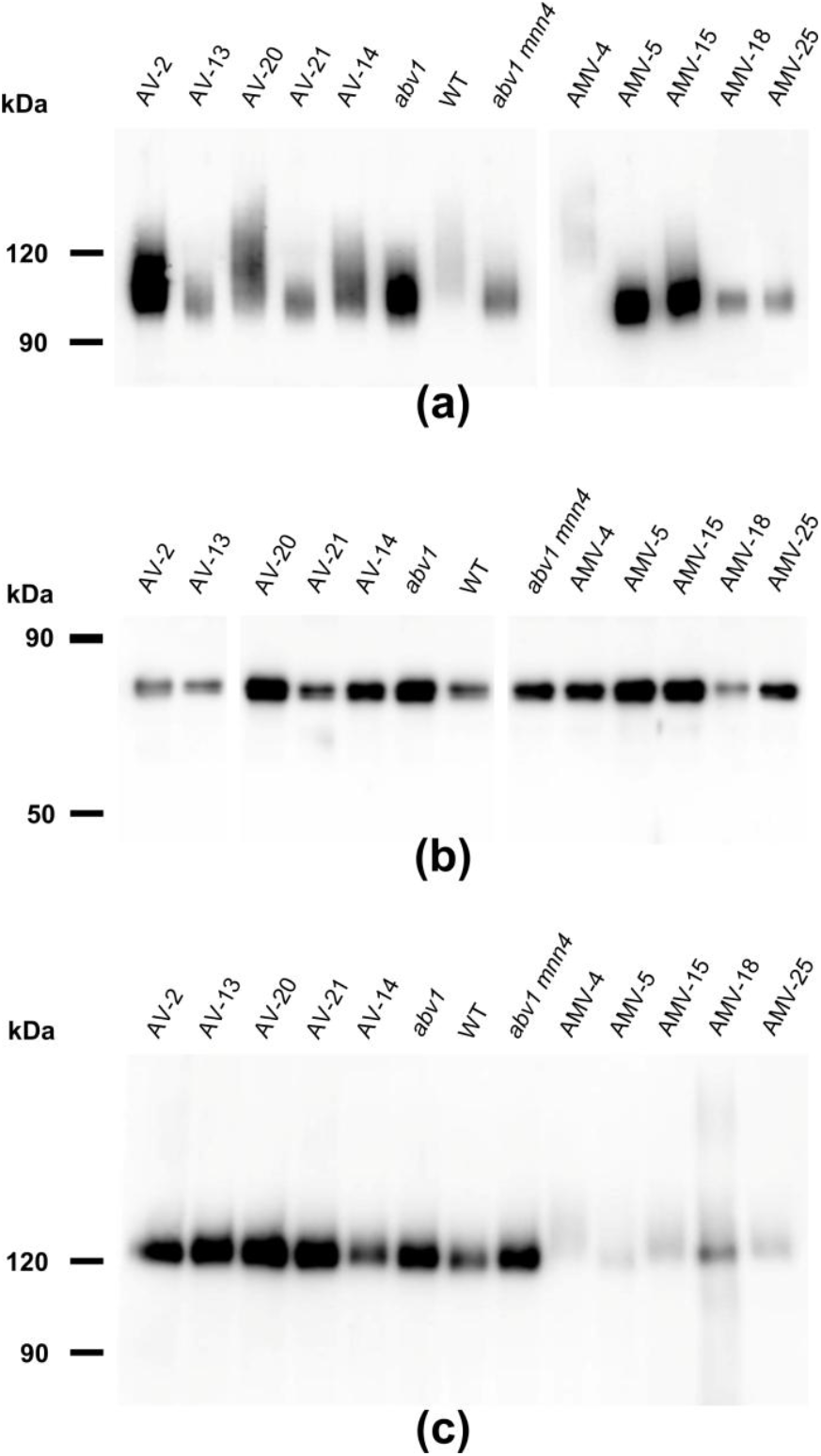
SDS-PAGE and immunoblotting of proteins from culture supernatants. (a) and (b), GOX before and after deglycosylation, respectively; (c), extracellular chitinase. AV2, AV13, AV20, AV21, and AV14, vanadate resistant mutants obtained in the M257-620M1 (*abv1-Δ*); AMV4, AMV5, AMV15, AMV18, and AMV25, vanadate resistant mutants obtained in the M257-620M1-MP9 (*abv1-Δ mnn4*); WT, the M257 strain.

### Inactivation of a *O. polymorpha* homolog of the Golgi apparatus α-1,2-mannosyltransferases increases vanadate resistance in the *abv1-Δ* mutant

Since the AV35 mutant was hypersensitive to SDS and elevated temperature, we suggested that these phenotypes and vanadate resistance are caused by the same mutation. To identify the gene defined by this mutation, AV35 was transformed with an *O. polymorpha* genomic library and SDS resistant transformants were selected. Since the genomic library was based on the AMIpSL1 vector, which has a propensity for genome integration, the obtained clones produced subclones with a mitotically stable plasmid. These subclones had decreased vanadate resistance compared to the original AV35 mutant. This indicated that the plasmids acquired by these transformants possess the wild-type gene, whose mutation led to vanadate resistance. These plasmids were recovered from yeast cells by the *E. coli* transformation and their sequencing revealed that they encoded a presumable α-1,2-mannosyltransferase [GenBank gene ID: XM_018356748.1 (Riley et al., 2016)]. Notably, *O. polymorpha* genome possesses two more open reading frames (GenBank ID: XP_018209282.1 and ORF locating at 1396245-1398152 positions of the scaffold NW_017264699.1) encoding proteins homologous to *S. cerevisiae* and *C. albicans* α-1,2-mannosyltransferases (Figure S2). Targeted disruption of the identified gene in the *abv1-Δ* mutant led to the same phenotypes as those revealed in the AV35 mutant. Inactivation of the identified gene in the strain with wild-type *ABV1* also increased sensitivities to SDS (Figure S3) and high temperature (Figure 2), but surprisingly, it caused a severe growth defect, which was not observed in the double mutant (Figure 3). This indicated that the *abv1-Δ* mutation may rescue the growth defect caused by inactivation of the identified gene. This gene was designated *ABV2* (*A*lcian *b*lue staining, *v*anadate resistance), since its mutations improved vanadate resistance in the *abv1-Δ* strain, while the cells of its knockout mutant obtained in the strain with wild-type *ABV1* did not bind the Alcian blue dye that indicated a severe defect of phosphomannosylation of cell-wall proteins (Figure 4).

**Figure 2.**
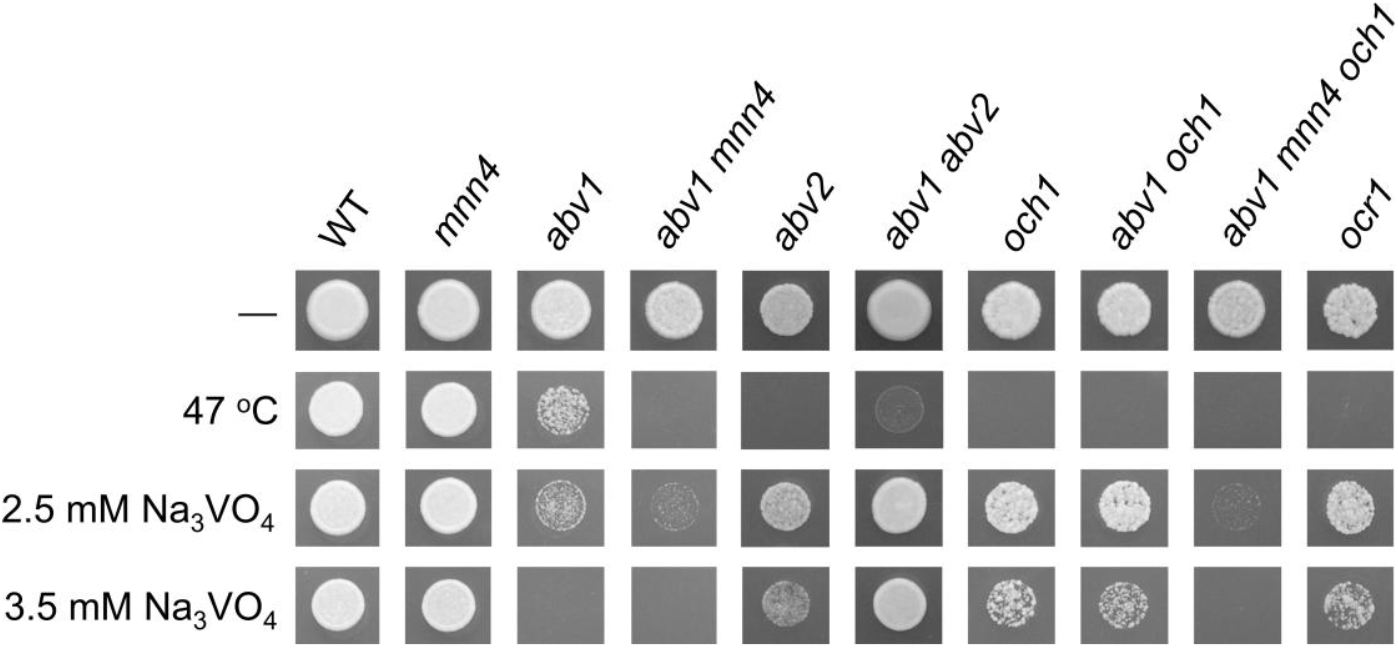
Sensitivities of M257-MP9 (*mnn4*), M257-620M1 (*abv1*), M257-620-MP9 (*abv1 mnn4*), M257-759M1 (*abv2*), M257-620-759M1 (*abv1 abv2*), M257-MP6 (*och1*), M257-1055 (*ocr1*), M257-620-MP6 (*abv1 och1*), and M257-620-MP9-MP6 (*abv1 mnn4 och1*) strains to elevated temperature and vanadate. The M257 strain was used as a wild type control (WT). Overnight cultures of all strains except M257-759M1 were 500-fold diluted with YPD medium and 4 μl aliquots were spotted onto the test plates. The culture of the M257-759M1 strain was 100-fold diluted due to its poor growth. The plates without vanadate were incubated overnight at 37°C (-) or 47°C (47°C), while the vanadate containing plates were incubated at 37°C for two days.

**Figure 3.**
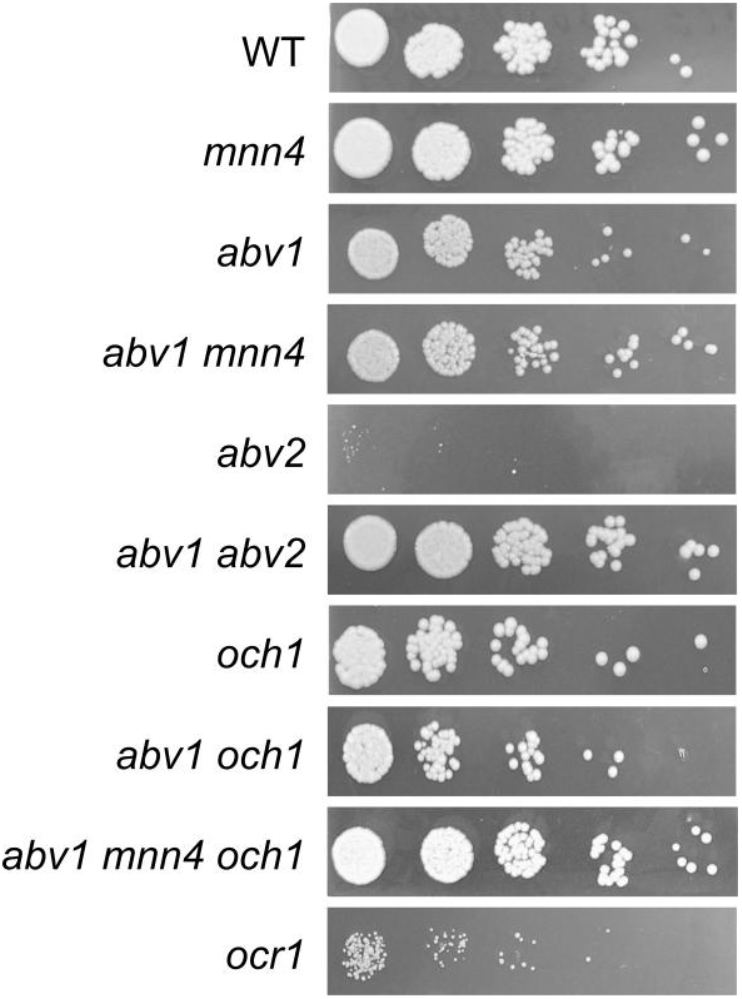
Growth of M257 (WT), M257-MP9 (*mnn4*), M257-620M1 (*abv1*), M257-620-MP9 (*abv1 mnn4*), M257-759M1 (*abv2*), M257-620-759M1 (*abv1 abv2*), M257-MP6 (*och1*), M257-620-MP6 (*abv1 och1*), M257-620-MP9-MP6 (*abv1 mnn4 och1*), and M257-1055 (*ocr1*) strains on solid YPD medium. Overnight YPD cultures were 1000-fold diluted with YPD and 4 additional 5-fold step dilutions were prepared and spotted onto YPD plate and incubated overnight at 37°C.

**Figure 4.**
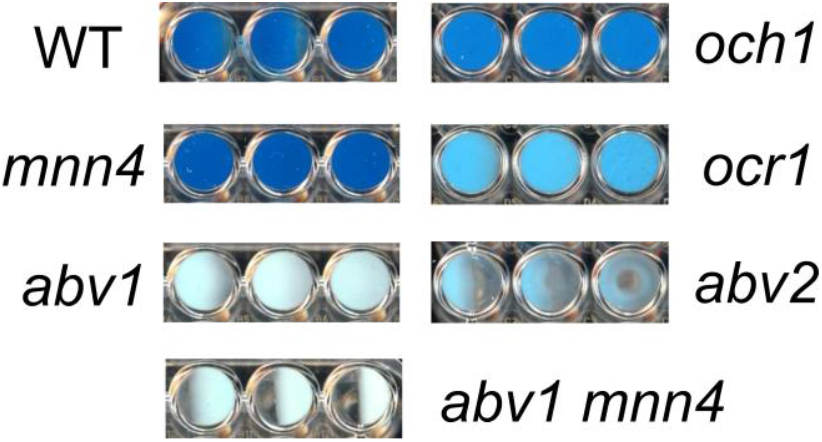
Alcian blue staining. WT, M257 strain; *mnn4*, M257-MP9 strain, *abv1*, M257-620M1 strain; *abv1 mnn4*, M257-620-MP9 strain; *abv2*, M257-759M1 strain; *och1*, M257-MP6 strain; *ocr1*, M257-1055 strain. Staining of cells from 3 independently obtained cultures is presented for each strain.

It was previously shown that deletion of the *O. parapolymorpha OCR1* gene encoding α-1,6-mannosyltransferase, which is responsible for the *N*-glycoside outer chain elongation, also causes a detergent and high temperature sensitivity as well as retardation in growth rate (Kim et al., 2006). These phenotypes were also revealed when we inactivated the *OCR1* gene in *O. polymorpha*, however the growth defect was less pronounced than in the *abv2-Δ* strain (Figure 3). Interestingly, inactivation of *OCR1* also led to a substantial decrease in Alcian blue staining, but it appeared to be not as severe as in the *abv2-Δ* mutant (Figure 4). Inactivation of the *OCH1* gene, which encodes another α-1,6-mannosyltransferase, also decreased Alcian blue staining, but less than the *ocr1-Δ* mutation. This correlated with the observation that in *O. parapolymorpha*, Och1 has a smaller effect on *N*-glycoside outer chain formation than Ocr1 (Kim et al., 2006).

### Inactivation of *MNN4* in the *abv1-Δ* mutant additionally decreases Alcian blue staining

Although some of the vanadate resistant mutants selected in the *abv1-Δ* mutant were defective in protein glycosylation in the secretory pathway, we could not exclude that phosphomannosylation was still involved in the increase in vanadate resistance in these mutants, since they possessed the *MNN4* gene, which encodes another homolog of the *S. cerevisiae* mannosylphosphate transferase Mnn4.

To ensure that the phosphomannosylation is completely abolished in *O. polymorpha*, inactivation of both *ABV1* and *MNN4* genes was required. However, we did not know whether inactivation of *MNN4* was lethal or not, how this mutation affects Alcian blue staining and how it interacted with *ABV1* inactivation. To explore this, the *MNN4* gene was inactivated in the *abv1-Δ* mutant and in the strain with wild-type *ABV1*. Both the single *mnn4-Δ* and the double *abv1-Δ mnn4-Δ* mutants were found to be viable. Inactivation of *MNN4* alone did not noticeably affect either vanadate resistance or Alcian blue staining (Figures 2 and 4). At the same time, inactivation of this gene in the *abv1-Δ* mutant additionally reduced Alcian blue staining (Figure, indicating that Mnn4 is involved in phosphomannosylation of a limited subset of the cell wall proteins or its mannosylphosphate transferase activity is much lower than that of Abv1.

Comparison of protein sequences of Mnn4 homologs from *S. cerevisiae, Yarrowia lipolytica, K. phaffii* and *O. polymorpha* (Figure 5) revealed that the common ancestor of these yeasts most probably possessed only one gene encoding a protein of this family, while the additional genes emerged after the lineages of *S. cerevisiae* and methylotrophic yeasts were separated. The *O. polymorpha* Mnn4 shows higher similarity to *K. phaffii* Pno1, while Abv1 falls into the same group with *Y. lipolytica* Mpo1 and *K. phaffii* Mnn4C and B. Alignment of protein sequences (Figure S4) revealed conservative domains present in all these proteins, while some positions distinguish the group comprising *O. polymorpha* Mnn4 and *K. phaffii* Pno1 from the group of the *O. polymorpha* Abv1, *K. phaffii* Mnn4C and *Y. lipolytica* Mpo1.

**Figure 5.**
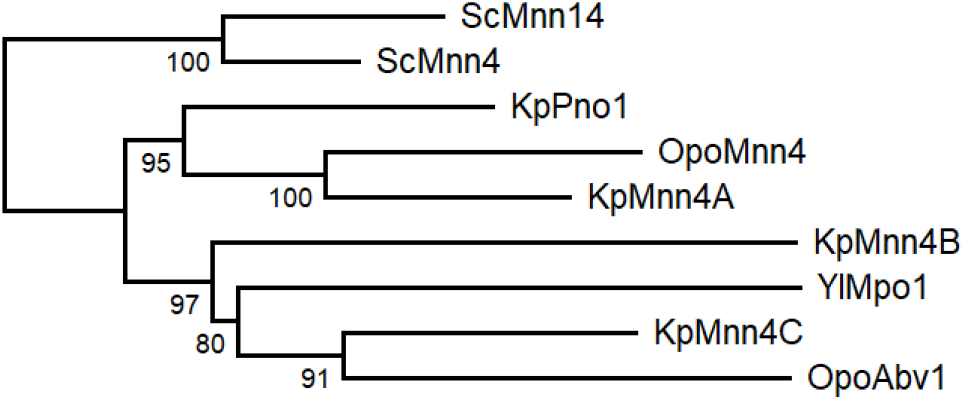
Phylogenetic analysis of yeast Mnn4 homologues from *S. cerevisiae* (Sc), *K. phaffii* (Kp), *O. polymorpha* (Opo) and *Y. lipolytica* (Yl). The Maximum Likelihood tree was created using the MAFFT (version 7.526) and the PhyML (version 3.0) program tools and edited using the MEGA (version 12.0.7).

### Mutations affecting *N*-linked glycosylation can increase vanadate resistance in *O. polymorpha abv1-Δ mnn4-Δ* double mutant

To obtain vanadate resistant mutants, the cells of the *abv1-Δ mnn4-Δ* strain were irradiated with UV and spread on plates with 3.5 mM sodium orthovanadate. Some of the obtained clones grew on regular YPD medium much slower than the original strain that complicated further analysis. The clones with acceptable growth rates on YPD were tested for sensitivity to SDS and to elevated temperature. Among 45 clones tested, 14 were unable to grow in presence of 0.004% SDS and at 46°C (class A), 11 were hypersensitive to SDS but not to elevated temperature (class B), 6 were hypersensitive only to the elevated temperature (class C), growth of 7 was similar to that of the original strain (class D), and 7 clones grew even better than the original strain in these stress conditions (class E). The latter class could arise due to suppression of the negative effects caused by *abv1-Δ* and *mnn4-Δ*. Nine A clones, 6 B clones, 4 C clones, 2 D clones and 3 E clones were examined for the GOX electrophoretic mobility (Figure S1, AMV mutants). GOX from some clones of class A and B, but not of other classes, demonstrated altered electrophoretic mobility indicating glycosylation defects. As described above for the mutants obtained in the *abv1-Δ* single deletion strain, the involvement of *N*-glycosylation in the difference in electrophoretic mobility was confirmed in some mutants by immunoblotting of glycosylated and deglycosylated GOX (Figure 1a and b). Presence of SDS sensitivity in the mutants with altered glycosylation was in agreement with previous observations that defects in protein glycosylation in the secretory pathway can cause detergent sensitivity in yeast (Kanik-Ennulat et al., 1995; Agaphonov et al., 2001; Kim et al., 2006). This phenotype is usually explained in terms of compromising the cell wall integrity and propensity of cells to lyse in the presence of a detergent. However other explanations can also be valid since we have shown previously that SDS interferes with Ca^2+^ homeostasis by promoting its uptake from the environment (Fokina et al., 2012; Kulakova et al., 2024).

Similar to the mutants obtained in the *abv1-Δ* strain, some of the mutants selected in the *abv1-Δ mnn4-Δ* strain secreted GOX with shorter *N*-glycoside chains than in the original strain (mutants AMV-5, AMV-18 and AMV-25), while in the mutant AMV-4 GOX glycosylation was significantly increased.

To study the effects of the obtained mutations on *O*-glycosylation, the electrophoretic mobility of extracellular chitinase was analyzed, since this protein possesses exclusively *O*-linked glycosides. Calculated molecular weight of its polypeptide chain is 59 kDa, but it migrates as an approximately 120 kDa protein due to *O*-glycosylation at multiple sites between the catalytic and binding domains (Kuranda and Robbins, 1991; Agaphonov et al., 2005). Five mutants obtained in the *abv1-Δ* strain and 5 mutants obtained in the *abv1-Δ mnn4-Δ* strain were studied. None of the mutants obtained in the *abv1-Δ* strain showed alterations in the chitinase electrophoretic mobility despite at least four of them (AV13, AV20, AV21, and AV14) having altered *N*-glycosylation of GOX (Figure 1c). The chitinase content in culture medium of these mutants was similar to that in the wild-type, *abv1-Δ*, and *abv1-Δ mnn4-Δ* strains. At the same time the mutants obtained in the *abv1-Δ mnn4-Δ* strain secreted noticeably less chitinase than the original strain and electrophoretic mobility of this protein was altered (Figure 1c). In particular, the AMV-4 mutant secreted more heavily glycosylated chitinase, while in AMV-5 it was less glycosylated. This correlated with alterations in GOX glycosylation in these two mutants. This means that these mutations affect the length of both *N*- and *O*-linked glycosides attached to proteins in the secretory pathway and indicates that selection of vanadate resistant clones in the *abv1-Δ*, and *abv1-Δ mnn4-Δ* strains may reveal different subsets of mutations affecting protein glycosylation. The latter suggestion was in agreement with the observation that inactivation of *OCH1* in the *abv1-Δ* strain slightly increased vanadate resistance, but had almost no effect in the *abv1-Δ mnn4-Δ* strain (Figure 2).

## Discussion

Previously, we have shown that vanadate resistance in the *Ogataea* yeasts depends on the *ABV1* gene expression level that affects regulation of the promoter of the *PHO84* gene encoding plasma membrane high-affinity phosphate transporter (Karginov et al., 2018). One could suggest that the presence of phosphomannose in the protein glycoside chains somehow downregulates the phosphate transport system and thus improves vanadate resistance. However here we show that even in the absence of the enzymes responsible for this modification other alterations of protein glycosylation can affect vanadate resistance in *O. polymorpha*. To do this, we first inactivated both *O. polymorpha* genes (*ABV1* and *MNN4*) encoding homologs of the *S. cerevisiae* Mnn4 enzyme, which catalyzes phosphomannosylation of glycoside chains, and then obtained mutations improving vanadate resistance and altering glycosylation of a reporter protein.

Unlike *ABV1*, inactivation of *MNN4*, which encodes another homolog of *S. cerevisiae* mannosylphosphate transferase Mnn4, did not noticeably affect vanadate resistance in *O. polymorpha*. Inactivation of *ABV1* alone conferred vanadate sensitivity sufficient to select genomic mutations improving vanadate resistance. One of these mutations defined a gene encoding a homologue of α-1,2-mannosyltransferases. Interestingly, inactivation of this gene in the strain with wild-type phosphomannosylation led to loss of the ability of cells to bind the Alcian blue dye. That was why this gene was designated *ABV2* (*A*lcian *b*lue staining, *V*anadate resistance), though the effect of its inactivation on vanadate resistance in the *abv1-Δ* mutant was opposite to the effect of *ABV1* inactivation in the wild-type strain.

Inactivation of *OCR1* encoding α-1,6-mannosyltransferase also led to the decrease in Alcian blue staining, however this effect was less pronounced than that in the *abv1-Δ* mutant. A noticeably weaker decrease in the dye binding was observed in response to the inactivation of *OCH1*, which encodes another α-1,6-mannosyltransferase. This correlated with the effects of these mutations on growth rate. We suggest that mannosylphosphate is attached to residues, which are attached by Abv2 to the *N-*glycoside outer chains initiated by Ocr1 or Och1. In this case the absence of either Och1 or Ocr1 alone should not completely abolish the outer chain formation in all *N*-linked glycosides and thus the glycosides, which have received the outer chain, can still be modified by the attachment of α-1,2-mannose and then by phosphomannose. At the same time Abv2 presumably can modify the outer chains independently of whether they were initiated by Och1 or Ocr1. Probably, that was why *ABV2* inactivation in the strain with the wild-type phosphomannosylation led to a more severe growth defect than inactivation of *OCR1* or *OCH1*.

Although the *O. polymorpha* Mnn4 protein shows higher similarity to *S. cerevisiae* Mnn4 than Abv1 does, the effect of its inactivation on Alcian blue staining is much less pronounced and no effect on vanadate resistance was observed. Possibly, this protein phosphomannosylates a specific subset of proteins, which are less represented in the cell wall and are not involved in the phosphate transport control. Interestingly, *Komagataella phaffii* has four Mnn4 homologs, namely Pno1 and Mnn4A-C (Miura et al., 2004; Bobrowicz et al., 2007). Inactivation of *PNO1* significantly reduced phosphomannosylation of a model recombinant glycoprotein, but did not noticeably affect Alcian blue staining (Miura et al., 2004). Since *O. polymorpha* Mnn4 shows higher similarity to *K. phaffii* Pno1 and Mnn4A, while Abv1 is more homologous to Mnn4B and C, it is reasonable to expect that the similarity of sequences represent similarity of functions and that *K. phaffii* Mnn4B and C should be more involved in phosphomannosylation of cell wall proteins than Pno1 and Mnn4A are. Some biotechnological applications require complete ablation of phosphomannosylation of recombinant proteins produced by yeast. This seems to be more easily achieved in *O. polymorpha* and *O. parapolymorpha* than in *K. phaffii*, since the former two species have only two genes responsible for this modification and as we have shown here both these genes can be inactivated simultaneously.

Selection of vanadate resistant clones appears to be an efficient approach to inactivation and identification of yeast genes responsible for the Golgi apparatus modifications of *N-* glycosides. To do this in *O. polymorpha*, here we first used the vanadate sensitive *abv1-Δ* mutant that allowed us to obtain mutants with altered glycosylation and identified the *ABV2* gene. The use of the *abv1-Δ mnn4-Δ* strain may allow obtaining mutations in different set of genes responsible for glycosylation. Indeed, the effect of *OCH1* inactivation on vanadate sensitivity was different between the *abv1-Δ* and *abv1-Δ mnn4-Δ* strains. The importance of *MNN4* for the manifestations of mutations affecting glycosylation is also highlighted by the observation that none of the tested vanadate resistant mutants selected in the *abv1-Δ* mutant showed alterations in *O*-glycosylation, while such alterations were revealed in mutants selected in the *abv1-Δ mnn4-Δ* strain.

## References

Agaphonov, M., and Alexandrov, A. (2014). Self-excising integrative yeast plasmid vectors containing an intronated recombinase gene. FEMS Yeast Res. 14, 1048–1054. doi: 10.1111/1567-1364.12197

Agaphonov, M. O. (2017). Improvement of a yeast self-excising integrative vector by prevention of expression leakage of the intronated Cre recombinase gene during plasmid maintenance in Escherichia coli. FEMS Microbiol. Lett. 364, 22–25. doi: 10.1093/femsle/fnx222

Agaphonov, M. O., Packeiser, A. N., Chechenova, M. B., Choi, E. S., and Ter-Avanesyan, M. D. (2001). Mutation of the homologue of GDP-mannose pyrophosphorylase alters cell wall structure, protein glycosylation and secretion in Hansenula polymorpha. Yeast 18, 391–402. doi: 10.1002/yea.678

Agaphonov, M. O., Sokolov, S. S., Romanova, N. V., Sohn, J. H., Kim, S. Y., Kalebina, T. S., et al. (2005). Mutation of the protein-O-mannosyltransferase enhances secretion of the human urokinase-type plasminogen activator in Hansenula polymorpha. Yeast 22, 1037–1047. doi: 10.1002/yea.1297

Aravind, L., and Koonin, E. V. (1999). The fukutin protein family - Predicted enzymes modifying cell-surface molecules. Curr. Biol. 9, 836–837. doi: 10.1016/s0960-9822(00)80039-1

Ballou, C. E. (1990). Isolation, characterization, and properties of Saccharomyces cerevisiae mnn mutants with nonconditional protein glycosylation defects. Methods Enzymol. 185, 440–70. Available at: http://www.ncbi.nlm.nih.gov/pubmed/2199792

Barone, G. D., Emmerstorfer-Augustin, A., Biundo, A., Pisano, I., Coccetti, P., Mapelli, V., et al. (2023). Industrial Production of Proteins with Pichia pastoris—Komagataella phaffii. Biomolecules 13, 441. doi: 10.3390/biom13030441

Bobrowicz, P., Terrance, S., and Stefan, W. (2007). Methods for eliminating mannosylphosphorylation of glycans in the production of glycoproteins.

Bowman, B. J. (1983). Vanadate uptake in Neurospora crassa occurs via phosphate transport system II. J. Bacteriol. 153, 286–291.

Bowman, B. J., Allen, K. E., and Slayman, C. W. (1983). Vanadate-resistant mutants of Neurospora crassa are deficient in a high-affinity phosphate transport system. J. Bacteriol. 153, 292–296.

Boyko, K. M., Varfolomeeva, L. A., Egorkin, N. A., Minyaev, M. E., Alekseeva, I. A., Favorskaya, A., et al. (2023). Preparation and Crystallographic Analysis of a Complex of SARS-CoV-2 S-Protein Receptor-Binding Domain with a Virus-Neutralizing Nanoantibody. Crystallogr. Reports 68, 864–871. doi: 10.1134/S1063774523601168

Bretthauer, R. K., and Castellino, F. J. (1999). Glycosylation of Pichia pastoris-derived proteins. Biotechnol. Appl. Biochem. 30, 193–200. doi: 10.1111/j.1470-8744.1999.tb00770.x

Choi, B.-K., Bobrowicz, P., Davidson, R. C., Hamilton, S. R., Kung, D. H., Li, H., et al. (2003). Use of combinatorial genetic libraries to humanize N-linked glycosylation in the yeast Pichia pastoris. Proc. Natl. Acad. Sci. 100, 5022–5027. doi: 10.1073/pnas.0931263100

Farofonova, V., Andreeva, N., Kulakovskaya, E., Karginov, A., Agaphonov, M., and Kulakovskaya, T. (2023). Multiple effects of the PHO91 gene knockout in Ogataea parapolymorpha. Folia Microbiol. (Praha). 68, 587–593. doi: 10.1007/s12223-023-01039-x

Farofonova, V., Karginov, A., Zvonarev, A., Kulakovskaya, E., Agaphonov, M., and Kulakovskaya, T. (2024). Inability of Ogataea parapolymorpha pho91-Δ mutant to produce active methanol oxidase can be compensated by inactivation of the PHO87 gene. Folia Microbiol. (Praha). doi: 10.1007/s12223-024-01236-2

Fokina, A. V., Sokolov, S. S., Kang, H. A., Kalebina, T. S., Ter-Avanesyan, M. D., and Agaphonov, M. O. (2012). Inactivation of Pmc1 vacuolar Ca2+ ATPase causes G2 cell cycle delay in Hansenula polymorpha. Cell Cycle 11, 778–784. doi: 10.4161/cc.11.4.19220

Giots, F., Donaton, M. C. V, and Thevelein, J. M. (2003). Inorganic phosphate is sensed by specific phosphate carriers and acts in concert with glucose as a nutrient signal for activation of the protein kinase A pathway in the yeast Saccharomyces cerevisiae. Mol. Microbiol. 47, 1163–81. Available at: http://www.ncbi.nlm.nih.gov/pubmed/12581367

Jigami, Y., and Odani, T. (1999). Mannosylphosphate transfer to yeast mannan. Biochim. Biophys. Acta - Gen. Subj. 1426, 335–345. doi: 10.1016/S0304-4165(98)00134-2

Kanik-Ennulat, C., Montalvo, E., and Neff, N. (1995). Sodium orthovanadate-resistant mutants of Saccharomyces cerevisiae show defects in Golgi-mediated protein glycosylation, sporulation and detergent resistance. Genetics 140, 933–43. Available at: http://www.ncbi.nlm.nih.gov/pubmed/7672592

Karginov, A. V., Alexandrov, A. I., Kushnirov, V. V., and Agaphonov, M. O. (2021). Perturbations in the heme and siroheme biosynthesis pathways causing accumulation of fluorescent free base porphyrins and auxotrophy in Ogataea yeasts. J. Fungi 7, 884. doi: 10.3390/jof7100884

Karginov, A. V., Fokina, A. V., Kang, H. A., Kalebina, T. S., Sabirzyanova, T. A., Ter-Avanesyan, M. D., et al. (2018). Dissection of differential vanadate sensitivity in two Ogataea species links protein glycosylation and phosphate transport regulation. Sci. Rep. 8, 16428. doi: 10.1038/s41598-018-34888-5

Kim, H., Moon, H. Y., Lee, D., Cheon, S. A., Yoo, S. J., Park, J.-N., et al. (2013). Functional and molecular characterization of novel Hansenula polymorpha genes, HpPMT5 and HpPMT6, encoding protein O-mannosyltransferases. Fungal Genet. Biol. 58–59, 10–24. doi: 10.1016/j.fgb.2013.08.003

Kim, M. W., Kim, E. J., Kim, J. Y., Park, J.-S. S., Oh, D.-B. B., Shimma, Y. I., et al. (2006). Functional characterization of the Hansenula polymorpha HOC1, OCH1, and OCR1 genes as members of the yeast OCH1 mannosyltransferase family involved in protein glycosylation. J. Biol. Chem. 281, 6261–6272. doi: 10.1074/jbc.M508507200

Kim, M. W., Rhee, S. K., Kim, J. Y., Shimma, Y. I., Chiba, Y., Jigami, Y., et al. (2004). Characterization of N-linked oligosaccharides assembled on secretory recombinant glucose oxidase and cell wall mannoproteins from the methylotrophic yeast Hansenula polymorpha. Glycobiology 14, 243–251. doi: 10.1093/glycob/cwh030

Kulakova, M., Pakhomova, M., Bidiuk, V., Ershov, A., Alexandrov, A., and Agaphonov, M. (2024). High-Affinity Plasma Membrane Ca2+ Channel Cch1 Modulates Adaptation to Sodium Dodecyl Sulfate-Triggered Rise in Cytosolic Ca2+ Concentration in Ogataea parapolymorpha. Int. J. Mol. Sci. 25, 11450. doi: 10.3390/ijms252111450

Kuranda, M. J., and Robbins, P. W. (1991). Chitinase is required for cell separation during growth of Saccharomyces cerevisiae. J. Biol. Chem. 266, 19758–67. Available at: http://www.ncbi.nlm.nih.gov/pubmed/1918080

Laemmli, U. K. (1970). Cleavage of structural proteins during the assembly of the head of bacteriophage T4. Nature 227, 680–5. Available at: http://www.ncbi.nlm.nih.gov/pubmed/5432063

Li, X., Shen, J., Chen, X., Chen, L., Wan, S., Qiu, X., et al. (2022). Humanization of Yeasts for Glycan-Type End-Products. Front. Microbiol. 13. doi: 10.3389/fmicb.2022.930658

Madhavan, A., Arun, K. B., Sindhu, R., Krishnamoorthy, J., Reshmy, R., Sirohi, R., et al. (2021). Customized yeast cell factories for biopharmaceuticals: from cell engineering to process scale up. Microb. Cell Fact. 20, 124. doi: 10.1186/s12934-021-01617-z

Mahanty, S. K., Khaware, R., Ansari, S., Gupta, P., and Prasad, R. (1991). Vanadate-resistant mutants of Candida albicans show alterations in phosphate uptake. FEMS Microbiol. Lett. 68, 163–6. Available at: http://www.ncbi.nlm.nih.gov/pubmed/1778439

Manfrão-Netto, J. H. C., Gomes, A. M. V., and Parachin, N. S. (2019). Advances in Using Hansenula polymorpha as Chassis for Recombinant Protein Production. Front. Bioeng. Biotechnol. 7. doi: 10.3389/fbioe.2019.00094

Mannazzu, I., Guerra, E., Strabbioli, R., Masia, A., Maestrale, G. B., Zoroddu, M. A., et al. (1997). Vanadium affects vacuolation and phosphate metabolism in Hansenula polymorpha. FEMS Microbiol. Lett. 147, 23–8. Available at: http://www.ncbi.nlm.nih.gov/pubmed/2038328

Miura, M., Hirose, M., Miwa, T., Kuwae, S., and Ohi, H. (2004). Cloning and characterization in Pichia pastoris of PNO1 gene required for phosphomannosylation of N-linked oligosaccharides. Gene 324, 129–137. doi: 10.1016/j.gene.2003.09.023

Munro, S. (2001). What can yeast tell us about N-linked glycosylation in the Golgi apparatus? FEBS Lett. 498, 223–227. doi: 10.1016/S0014-5793(01)02488-7

Ravin, N. V, Eldarov, M. A., Kadnikov, V. V, Beletsky, A. V, Schneider, J., Mardanova, E. S., et al. (2013). Genome sequence and analysis of methylotrophic yeast Hansenula polymorpha DL1. BMC Genomics 14, 837. doi: 10.1186/1471-2164-14-837

Riley, R., Haridas, S., Wolfe, K. H., Lopes, M. R., Hittinger, C. T., Göker, M., et al. (2016). Comparative genomics of biotechnologically important yeasts. Proc. Natl. Acad. Sci. U. S. A. 113, 9882–7. doi: 10.1073/pnas.1603941113

Sohn, J. H., Choi, E. S., Kim, C. H., Agaphonov, M. O., Ter-Avanesyan, M. D., Rhee, J. S., et al. (1996). A novel autonomously replicating sequence (ARS) for multiple integration in the yeast Hansenula polymorpha DL-1. J. Bacteriol. 178, 4420–4428. Available at: http://www.ncbi.nlm.nih.gov/pubmed/8755868

Stankiewicz, P. J., Tracey, A. S., and Crans, D. C. (1995). Inhibition of phosphate-metabolizing enzymes by oxovanadium(V) complexes. Met. Ions Biol. Syst. 31, 287–324. Available at: http://www.ncbi.nlm.nih.gov/pubmed/8564811

Towbin, H., Staehelin, T., and Gordon, J. (1979). Electrophoretic transfer of proteins from polyacrylamide gels to nitrocellulose sheets: procedure and some applications. Proc. Natl. Acad. Sci. U. S. A. 76, 4350–4354. doi: 10.1073/pnas.76.9.4350

Veale, R. A., Giuseppin, M. L. F., Van Eijk, H. M. J., Sudbery, P. E., and Verrips, C. T. (1992). Development of a strain of Hansenula polymorpha for the efficient expression of guar α-galactosidase. Yeast 8, 361–372. doi: 10.1002/yea.320080504

