## Supplemental figures and table for "Alterations in protein *N*-glycosylation confer vanadate resistance in *Ogataea polymorpha* mutants defective in phosphomannosylation"

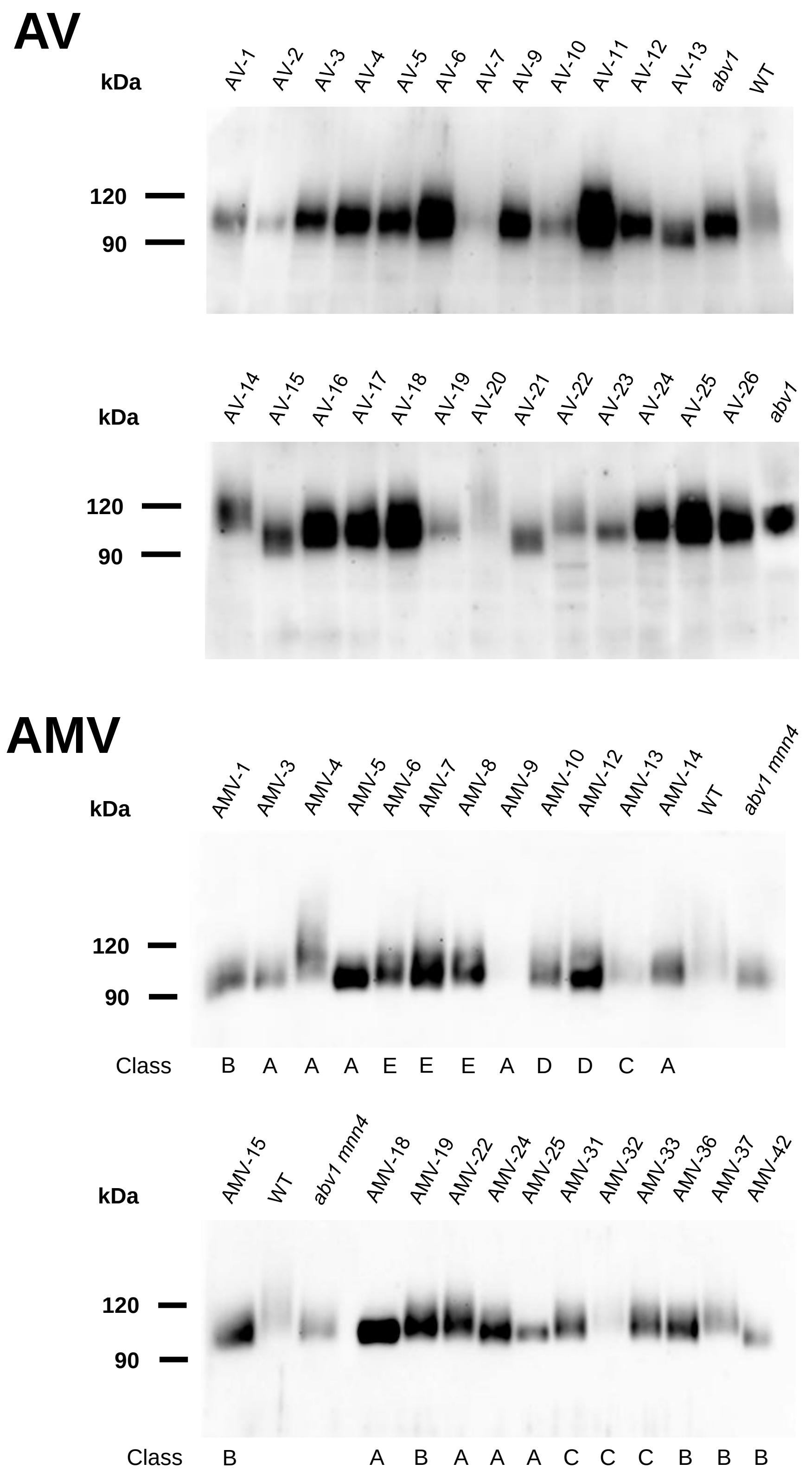


Figure S1. SDS-PAGE and immunoblotting of extracellular GOX. AV, culture supernatants of mutants obtained in the *abv1* mutant M257-620M1, whose culture medium was used as a control (*abv1*). AMV, culture supernatants of mutants obtained in the *abv1 mnn4* mutant M257-620-MP6, whose culture medium was used as a control (*abv1 mnn4*). Culture medium of the M257 strain with wild-type glycosylation (WT) was used for comparison. Classes of the AMV mutants are indicated at the bottom: class A, clones exhibiting sensitivity to both elevated temperature and SDS; B, clones sensitive to SDS only; C, clones sensitive to to elevated temperature only; D, clones growing in these stress conditions similar to the original strain; E, growing better that the original strain.

```
          1          10        20        30
OpoAbv2   MS........MLFARNRRAK...TLILVVSILITVLVLSSS..............AGTEH
OpoMnn2A  M............................................................
OpoMnn2B  MW...ITKLNLKYRTKQRVRI..SLILACAAFMLLFLVVRSFFLE..NDERV........
ScMnn2    M.........LLTKRFSKLF...KLTFIVLILCGLFVITNK...................
ScMnn5    M.........LIRLKKRKIL...QVIVSAVVLILFFCSVH....................
CaMnn2    MI............AKQKIK...ILIGVIIVIATYHFIVSSNVRS...............
CaMnn21   MFQQLTYRLRLFRRRHKYIFI.NSIFLSVIIIFLIYSYWSNLPAE..DNSAIINEKGTYH
CaMnn22   MG........SIFKDGRRILVRPKSL..IICLCLISIIFTQL..................
CaMnn23   MS........INFLSIPRNRFKAIGVLSVTCILIYVILHSSI..................
CaMnn24   MF........SIPVSSKTVRLILVSLLLITLINILAAFQRSTLSSWFPSSRHIINKFTDL
CaMnn26   MS........LRRLSPSHLIL.GTLVLGVIIFNLYVLTSTHE.....DIKKV...KGPTY

             40        50        60        70        80
OpoAbv2   LSKFKLPTSSSYSIYAASEVKEEPSGSAAPSSTPEDAENLRNLYFSPEQGGV........
OpoMnn2A  ............................................................
OpoMnn2B  ............................................................
ScMnn2    ............................................................
ScMnn5    ............................................................
CaMnn2    ............................................................
CaMnn21   RSLWESITMALFP......PKTKPFEEKKPQVNPNNNQEV..........GV........
CaMnn22   ............................................................
CaMnn23   ............................................................
CaMnn24   RLALSSQESVLRD......EEGEIYSLVGYHHDFDS.....NLVVVQKQ...........
CaMnn26   HTSDNTKIQSHISNYDSEEYVDRLTAEIEDAKKEELISEIRKKLEIQEKPGVIQKLKTEL

            90                         100         110
OpoAbv2   .IPG..................GSSGNTKGWKVSSIDESK...GQRTD...........
OpoMnn2A  ............................................................
OpoMnn2B  ..AGSSAIVVNNDLAQR......PTSKKTAKELLAQKKKTR...ESVYD...........
ScMnn2    ..................YMDENTSVKEYKEYLDRYVQSY...SNKYSSSS........
ScMnn5    ....................NDVSSSWLYGKKLRLPVLTR...SN..............
CaMnn2    ..KDLSDLV..........DLGSSDKSTTENERPKNNIVT...NNRLDNP.........
CaMnn21   .ESGASEISQHKQQQQQQQHAKEPTTKTSSKSLVSD........EAYQL..........
CaMnn22   ......................IRYQYQL..IADEVQPTI...NEDHSSS.........
CaMnn23   ......................ITTDFDVSDYGDKFIPSII.FDDNNDNG.........
CaMnn24   ..................YLLEKTPNEDTTEHFWNFLQSNFETKSEYDLNLI.......
CaMnn26   RMKYIDDIKN.........HLKQEITEQYSNEIFKQYAFSFEIYSKKVDEYALQLENSLK

                          120        130       140       150
OpoAbv2   .............DRSIPLTENQ..SPTEEALNKITPEDIVRGDVTF.QNFFIEIF.EL
OpoMnn2A  ...............................................:....PL.EV
OpoMnn2B  .............NLSVEDREKM......RVINQPSPDDPRGLRLELGHTFFKSVF.NV
ScMnn2    .........DAASADDSTPLRDND..EAGNEKL...............KSFYNNVF.NF
ScMnn5    ..................LKNN..........................FYTTLV.QA
CaMnn2    ..........PNEDIPHAEPDSP...........PQEPPKSGNKPDFSIFFEGL..EK
CaMnn21   .............NLKLIN.EQQ......EKYNRLHPFEQK..RLQQGYEFFDNIF.KI
CaMnn22   ..........QSLKNTK..LNST..RSSSPI......SPPKSLNKLTSQEFWEHIF.NI
CaMnn23   ..........ENLKDPQPELDND..KGNGET......DTTTSNSMTTAHTFWKGIF.DT
CaMnn24   .........DGYNYKKLIKHLNEQ..NELQLSHSFVEQYKMENQFIQSFQNFFVQLI.DT
CaMnn26   PASCLAILQQAEKDTDSIPDLENYLTKANDKYFKR..............QEYWRYLLKDI

                   160           170                             180
OpoAbv2   MRK........NQLSYPL.....AERQTLKD..................G.KP..FIEN
OpoMnn2A  LVK........REYGAYK.....AVNVAMHDTV................G.VP..L...
OpoMnn2B  LDKGKPRVSPLTKYKSTE.....RIYHAGYEVM................GDGP..S...
ScMnn2    LMVDSPKGSTAKQYN.........EACLLKGDI................GDRP..DHYK
ScMnn5    IVENKPADSSPDLSKLHG.....AEGCSFANNV................AAHD..SGHD
CaMnn2    FAIKQPGIK..DKYTSEK.....AKEKFSTDDN..................:......
CaMnn21   FYQAKPSVSQLNTYPSKK.....RIYHARFDSL................ADDD..T...
CaMnn22   FEINKF........................DD.................GLKP..LIKY
CaMnn23   FDKYKM........................DL.................GQDPENAVSY
CaMnn24   IEDCKPDLDPINNDNHYPNGDKIVKKYYELRNKIPSENMKQFNIERLIHRNGRIP..IYGG
CaMnn26   LLNNKPKCEPLTKEEKGE...............................KLNP..TYQW

                190          200         210        220
OpoAbv2   VLFFAQPWDRL.SEED.CLQFVNFPPQFINDLTLKHETVV.SNLPK..............
OpoMnn2A  ..........L.SEEY.LLNCLKVPPEMVADLAVSHRNVV.RDLPA..............
OpoMnn2B  ..........L.SKEY.LSRFLQLSRSEINSMKKSHSFVT.RNLPE..............
ScMnn2    DLY......KL.SAKE.LSKCLELSPDEVASLTKSHKDYV.EHIA.............TL
ScMnn5    S........DL.SYES.LSKCYNLNKTVQESLREVHSKFT.DTLSGKLN........FSI
CaMnn2    .........FLFGKEY.LENVLDIPQATFKELKDSHKRYVDEHIPKMLQRVKTFGSLAPS
CaMnn21   ..........IFSEKY.LSQFLQLSNEELAAMKKSHKYVV.ENLPE..............
CaMnn22   TPKEQQLTTKIKTRDW.LLSKANIFHKDI..IRQYHESVL.RNLPS..............
CaMnn23   VDKLKQKQGPL.NKEV.LLSKAIVSSELMKHLKEKHSGVV.EDLPS..............
CaMnn24   HLREQYKDELIRNKEF.LSMYLTLSDSEISALKKSHTKFL.ETMME..............
CaMnn26   DAR......II.SEQYLLGSKLTIPGEKFRALRSAHDQVV.KQLKS.LP...........
```

```
                 230                240       250       260       270
OpoAbv2     ITPR.FYKGN.........GYVIVGGGKYSWFALLGIETLRKVGSTLPVEVILPSDDEY.
OpoMnn2A    SYPEGLYEGR.........GIVFVGGGKFSWLSLLGIENLRATGSKLPVELIFPTEAEY.
OpoMnn2B    TYPIGLYSGN.........GVVYVGGGKFNWLALLSIKTLRSVGSKLPVEILIPKLDEY.
ScMnn2      VSPKGTYKGS.........GIATVGGGKFSLMAFLIIKTLRNMGTTLPVEVLIPPGDEG.
ScMnn5      PQREALFSGS........EGIVTIGGGKYSVLAYTMIKKLRDTGTTLPIEVIIPPQDEG.
CaMnn2      DKEWESYKGS........SGYIIVGGGRFTWLSFLVIKQLRATGAKLPVEMFIATESDY.
CaMnn21     DAPDGLYKKN.........GIVYVAGGSFNWLTLLSIKSLRAVGCHLPIEVFIPKIEEY.
CaMnn22     KLPKSVYTPNT.......YGIVTIGGNFYSWMAYIQLLQLRKLGSNLPVEILIPSIEDYY
CaMnn23     VMPGSVYNKGS.......KGVVIIGGGKFSWLAYLALVQLRNVGSKLPVEIVMPSRADYE
CaMnn24     NWPENLFKFNKFNNFMKGDGIVYLGGGKYNQLVLLSIKILRENGSRLPVEVIIPYKNDY.
CaMnn26     DPPSQFISGH.........GIVVNGGGNMIGSALTAIANMRERGSQLPVELILDTKQEY.

                  280       290               300       310       320
OpoAbv2     .EFEYCDQILP.ALNARCVEMPRVF.......GKTTLRKFDVNGYQFKAFALFASTFENA
OpoMnn2A    .EEMLCEKVLP.DLNAKCVLLTERV.......PEFRKHKYTIRGYQYKILALLVSSFEQV
OpoMnn2B    .EVDLCTTIFP.ALNAKCIYMPKQL.......GEQISERFSFFGYQYKALALMLSSFENV
ScMnn2      .ETEFCNKILP.KYNSKCIYVSDIL.......PRETIEKFVFKGYQFKSLALIASSFENL
ScMnn5      .EDDFCKNWLP.KFNGKCIYFSDIV.......PSKPLSDLKLTHFQLKVFGLIISSFKRI
CaMnn2      .EKEFCEKVLP.KYNARCNVFDYKL.......ADDLKKRFDIGGYQYKMLALLSSKFENV
CaMnn21     .ESDLCNRILP.ELDARCIYMKNQLMNPNKDNSDSFANKFEFKGYQYKALAILLSSFENV
CaMnn22     KEAHFCDHVLP.QYNAKCILVPEKL.......GFNVAKHWKFSSYQFKALALCLSSFQHV
CaMnn23     KELEFCENTLP.EMQASCVVLPDVL.......GEAVMKNRKFASYQFKALALVVTSFEHI
CaMnn24     .DIQFCDRVLP.TLNGKCKLMTDYL.......PQTFVDK..ISGFQLKNIALLISSFERI
CaMnn26     .DKQICEELLPKKLNGKCVIVEEQV.......GKEVFD.IINEKFSRKIMGLLVSSFDHI

                330       340       350       360       370        380
OpoAbv2     FFLDSDAYPVANPDPLFESDLYKEYQMITWPDFWRRTTSPYFYQITGQEIGPKQ.VRHLN
OpoMnn2A    LFLDSDNVPVANPDAIFVSEPFTSHGMVTWPDFWRRVTHPTYYKVVDRELGNEQ.VRNNI
OpoMnn2B    LLLDADNTPLHAPDHLFETEPFTSTGMVIWPDYWKRSTSPAFYDIVNIEIDEGH.RVSHG
ScMnn2      LLLDADNFPIKPLDNIFNEEPYVSTGLVMWPDFWRRTTHPLYYDIAGIAVDKKKRVRNSR
ScMnn5      IFLDADNYAVKNLDLAFNTTSFNDTGLILWPDFWRRVTPPAFYNIIGSSINIGKRVRFVS
CaMnn2      LYLDSDNFPTRNVDYLFESDLYKENNLLLWPDAWARTTNPKYYEIAGVPVKENK.LRYSK
CaMnn21     LLLDSDNIPAHSPEELFENDPFKSYGLVVWPDYWKRATSPYYYNIADIDVSEKY.LGSKY
CaMnn22     LILDSDNVVLSKPEKVFDSPVYRDNGMVLWPDYWERTISPEWYDIIGKPVVGNKQVRTGR
CaMnn23     LLLDSDNMIVSNPDEIFESKLYHQYGMITWPDYWKRTISPLFYDVAEIEVNENKRVRYNR
CaMnn24     LYLDADNIPIRNPDVLFTNAPFTTKHLVVWPDLWRRSTSPHYYTIAGIEVDPNFKVRNSY
CaMnn26     IAMDADNLAIKNVDNLLFTEPYLSTKMILWPDLWVKLTSPLYYKIARIEPGEIV.DRFGI

                  390                      400       410       420
OpoAbv2     DMF.TDPKYYE....SEL...........NADPYHNIPFHDREGTIPDWTTEAGEMLINK
OpoMnn2A    DDV.TPNKYYA....RD............TGHAFSSMPLHDRAGALPDPSSESGQIAVDK
OpoMnn2B    FQE.YGK..YT....TP............NSPPDQAPPLHQLKGAIPDPSSESGQLMLSK
ScMnn2      DDI.TPPAVYT....KDLKDLSD............VPLSDLDGTIPDVSTESGQLMINK
ScMnn5      DDI.SPVSRYD....PFVSNSNDYTPKERQEHFLKHVPLHDLDGTMPDLSSESGQMVIDK
CaMnn2      YDEKQAGGKDK...LKPL...........SEYTFKDSWYHDFEGTLPDPTSETGMFMVNK
CaMnn21     NEV.EGQ..YTDLSVEK............GSVELDKIPLHQRLGSIPDPTSESGQLLISK
CaMnn22     FPV....NIHN....MLT...........SELEINETRFHDLEGALPDLSTESGQVMFNK
CaMnn23     FPLYNAPNVRS....NIY...........TDQEREEVPFHDLQGSIAELSTESGQLIINK
CaMnn24     VDG.DERGKYT....DSM...............YYSYHDCKGSIPEASSETGQLLINK
CaMnn26     PND....ASFA....EYI...........TKDKQSEVHYHDLDNLPSTISVETGQMVFSK

              430       440       450       460      470       480
OpoAbv2     TLHFQTLLLALYYNFDGPYGYYPLLSQGGAGEGDKETFVAAANYYGLKYYQVYKLPDRAY
OpoMnn2A    RTHLRALLLALYYNYYGPQQYYPLFSQGGAGEGDKETFVAAAQHFGLPFYHVRKAVDVIG
OpoMnn2B    KTHSKVMLLALYYNMYGPNHYYPLLSQGSDGEGDKETFIAAAHVLKKSFYQVKKIIKSIG
ScMnn2      TKHLATALLSLFYNVNGPTWYYPIFSQKAAGEGDKETFIAAANFYGLSFYQVRTRTGVEG
ScMnn5      IRHFNTLLLALYYNVYGPTWYYKMISQGTAGEGDKDTFVAAAHALNMPYYQVRTNFEFDG
CaMnn2      SSHLKTLLLCLYYNVFGPQYYYPLLTQGSAGEGDKETFIAAAHVMKEPWYQCARQFKWTG
CaMnn21     KTHLKPLLLALYYNLYGPSHYYPLFSQGSDGEGDKETFLAATVTLGKRYYQVAKFLVSLG
CaMnn22     KTHGKVMLMTLYYNIFGPEIYYKLFSLGALGEGDKDTFAAAALACGEKYYQVASSIRTLG
CaMnn23     HTHGKTILLALYYNFYGPNLFYKLFSLGEQGEGDKDTFVAAAVVTRQDYYQVKSFIKTFG
CaMnn24     KIHFQTLILAMYYNYYGPDYYYPLFSQGAAGEGDKETFIAAAHKLDLPYYQVGEFNREFG
CaMnn26     REHLKSLLLALYYNINGKDFYIDLLYQGAYGEGDRETIVPALHVMNERYSLTNHKVHILG

               490       500       510                           520
OpoAbv2     GW.YNHEQNYEHSSIVQYDPLTDYSNL........QQ......VKQNI.......RKAIE
OpoMnn2A    YWQLQPEEHYTGVGMIQYDPVVDYKLVAAYKDWFSAR......EKEHE.......RRQAS
OpoMnn2B    RWA...NDEFTGSAMGQFNCHEDYKLYK...............................K
ScMnn2      .Y..HDEDGFHGVAMLQHDFVQDYGRY........LN......AMESIGNKY.GGTKSAD
ScMnn5      FF..YQKDDYKGLALLQHDFEQDYKQY........QK......AQQKV......KANIEE
CaMnn2      YVS.KVDNKFTSKALAHYDPVQ.....................AQDTT............
CaMnn21     HFKV.PGGDFEGCGMGQFDPQQDLEYIKLREQYAKIP......EKDKE........KQHK
CaMnn22     YFDTTPGGGFHGMAMAQKNPQLDYQLF........QK......TNQNF........KDLH
CaMnn23     YADSD..DKFQGVSMGQRNPLIDRKHY........EDHVLALLEKDSF........KSSS
CaMnn24     PIN.DNTRKHEFYGMGQYDPIIDYYMSTI......TT......TQKDT........KKKIN
CaMnn26     YD..APNGKYSETTLGQTDPRDGFEFYQDWRKFLTSR......KLDTRLNPFQSGGYTSD
```

```
                         530          540                   550          560
OpoAbv2    LEGE.......DFKYDYEK.FFTQFFL..........PSNSKVLFYHVHDPKMDPFNIV.
OpoMnn2A   FIGRLRDWWAGDVVYDPAR.TFTGWFN..........KTNSRPMFVHSNFPKLDPVSLR.
OpoMnn2B   YENSFE..................HV..........DEVPKLLFLHSNFPKLDPWSLK.
ScMnn2     AIKF......DKNYSLEK.YTEEFFDNEDLN........AKNHVDVMFIHSNFPKFDPYDLS.
ScMnn5     FSKL......DPDYTLDNGFLKTLMVNDD.......GSDLDIMFIHASFYKADPWTLY.
CaMnn2     ....................................RQDVDIIFMHLSYPKFYPNWLA.
CaMnn21    FKLQHQ..................VL.........EKGPEILFVHANFPKLNPWKLK.
CaMnn22    L...............D......WDNEAKDQF.SANNKIPIFTMHCNIKKINPAAYM.
CaMnn23    IA......DQIEQMKK.....FENEDFDQ....HNSIPLFTVHCNYPKLDPKLYM.
CaMnn24    YNSPL......PEKYAAND..EDDTCSNYDFHL....FQSSSLFFLHANWPKYYIEKLFL
CaMnn26    LMKQFHDYK..RQIYQDKQ......YEDEAAVHRMITYKLPSILFLHCNHPKIDPLKNS.

                     570           580                                  590
OpoAbv2    ....NNRYTFDLD.G...NRIRNLGE.D......................F.PEF..DF
OpoMnn2A   ....KDDLLFVTN.GGARKRVR.MYS.D......................Q.SGL..DF
OpoMnn2B   ....VDEVIYDPETG...KRERLYGT.E......................YLEAL..GF
ScMnn2     ....KSN.FLTTN.G...KPARSYTALK......................KVKNY....
ScMnn5     ....HENRFIGPN.G...EQVRGF...R......................KPHRY..GM
CaMnn2     ....DNHDLVYADTD...EHIRMYSS........................INKNV..GY
CaMnn21    ....QEKKIFDAK.G...NRVRLYGP.G......................MIKRI..GY
CaMnn22    ....KDEKIANMD.E.KRMNVRFYSNLKFKLNDDDIYSPDNEKEKNKSEKTDNDPN..EI
CaMnn23    ....SRDDLYDEK.E.KKLKYRLYGNLK.......YTKEVVKDGETGTEASTDKV..QI
CaMnn24    YSYDEDRGPVTND.G...DKRRLYGN.E......................LKKELG..GY
CaMnn26    ....KEA...DAEFG..VYSRRNMGLPD......................KVEKLLEGK

                600         610                620          630          640
OpoAbv2    DLENFIWNVINHYVCEKKIN....FKHFEN....ADWNSL..CDRFIQDQLQFLAESSRR
OpoMnn2A   DYELRQWQFIKKHFCDKAID....LNYLRV....AGVTSLDYCD.FIEQELEFL.ENSQH
OpoMnn2B   DFERVQWENMKFYICDLNLH....PKLFSD....RDIDHDELCT.EIKEHLEFL.KSTTN
ScMnn2     DIELENFKVLNEYVCVNKNP....FKYLDDLLGQDKTEWKRVCG.YITDRLAFL.ESTHD
ScMnn5     DFELFLFNDMRGSFCTTPKSQVIKFKYFTDKV..NTPEWDAMCE.YLTNHVNYL.ESTHK
CaMnn2     DFDLRVMQFFTEGLCPNYYD....SKTGKP...IENIPHIDYTNDYMGNHLMYV.KDEEQ
CaMnn21    DFEWFQWEGMKYLLCIYDVN....LQIF......QDVKKSDLCD.EILEHLRFL.ESTQN
CaMnn22    DFELSRWQIVSEILCDQKIT....FQFLKD....ENM..DEVCQ.FVKNTIAWL.GK.KT
CaMnn23    DFELQQWQHMQDILCLKKIY....FTHFID....NDM..NELCQ.FIENQVAWL.SKSQN
CaMnn24    DFELNIMKNLHWCFCEEPLIDLIGIPVVGSKT......RTDVCI.AIKNHIEFL.EIS..
CaMnn26    DWELRFHTISQWVACEAISK....SSLYWEKI..AGKSQQQVCE.SVGKYIEFL.KKDTF

                     650         660                        670
OpoAbv2    .IFSDHNKSPPPDLVESPEKAEKVE.........TPDDV......AVE
OpoMnn2A   .LVNIRNATT......................................
OpoMnn2B   ................................................
ScMnn2     KAIAGK..........................................
ScMnn5     EAMGEKN.........................................
CaMnn2     NNIDRCNKVFIPHLKWLKETAEI..............PT......VVS
CaMnn21    .ITD............................................
CaMnn22    ................................................
CaMnn23    ................................................
CaMnn24    ................................................
CaMnn26    DNEATK..LTILNQLGEKSQPKQPEINNNNNNNNNDDDNGKNKQGAAS
```

Figure S2. Multiple alignment of amino acid sequences of Mnn2 homologs from *O. polymorpha* (Opo), *S. cerevisiae* (Sc), *Candida albicans* (Ca), generated using MAFFT and visualized in ESPript with BLOSUM62 matrix. Red background indicates strictly conserved residues; blue boxes highlight position with physicochemical similarity. OpoMnn2A, the polypeptide encoded by the ORF locating at 1396245-1398152 positions of the scaffold NW_017264699.1; OpoMnn2B, XP_018209282.1.

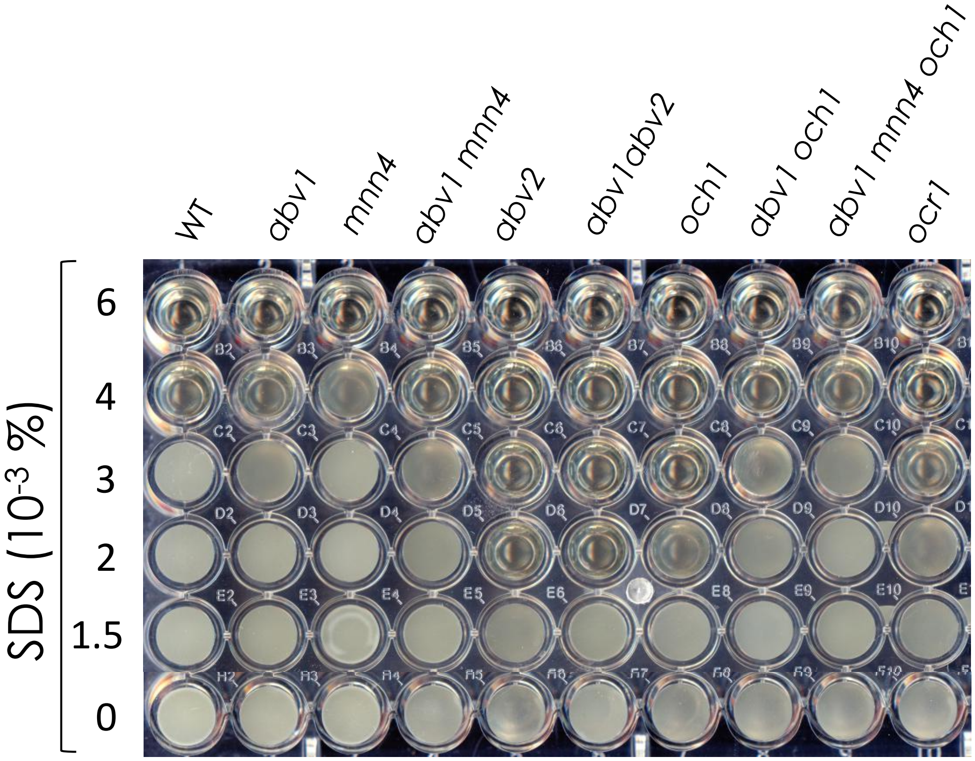


Figure S3. Growth of M257 (WT), M257-620M1 (*abv1*), M257-MP9 (*mnn4*), M257-620-MP9 (*abv1 mnn4*), M257-759M1 (*abv2*), M257-620-759M1 (*abv1 abv2*), M257-MP6 (*och1*), M257-620-MP6 (*abv1 och1*), M257-620-MP9-MP6 (*abv1 mnn4 och1*) and M257-1055 (*ocr1*) strains in liquid YPD supplemented with SDS in different concentrations.

```
          1
KpPno1    MT...................LRSAIKA................................
KpMn4A    MK..........................................................
OpoMnn4   MT....................SASNKR................................
ScMn4     ML...................QRISSK.................................
ScMnn14   MM..........................................................
YlMpo1    MV..........................................................
KpMn4C    MS.GNPFLFSPSNFDFSGLDHYRSTDKDHLALDVLDYDKNHFFSRNSPSLKSRIHFYRHK
OpoAbv1   MAQGDEELFIQ........EPPRAARKESHAS.........FILKAWLALVAPFTFV...
KpMn4B    MF....................KETSKNLFGS............................

                       10        20        30              40
KpPno1    ..............RTSKGLIGAVIIASIIFFTTVTF......YDESKIVGIIR..VSDT
KpMn4A    ......VSKRLIPRRSRLLIMMMLLVVYQLVVLVLGLES....VSEGKLASLLD..LGDW
OpoMnn4   ......YAVMMVPRRRRCVTLIAAICLLNLVIVSF........LGAGTQMDVLNDVTGLI
ScMn4     LH.RRFLSGLLRVKHYPLRRILLPLILLQIIIIITFIWSNSPQRNGLGRDADYLL.....
ScMnn14   LSLRRFSMYVLRSLRLHFKKIIITLLTIQLLFITIFV.......LGGRSSII........
YlMpo1    LH......PFRLLRTSLVSKLVVILITCLIFGSL..........................
KpMn4C    LT.TRKQIGLFSGRLKLFVLALFVLITFSAIHIPIPF.......................
OpoAbv1   LN.KFIQLLPYHNNRKMSVFTIYIGLMLVFIVPTIFYMDESI..................
KpMn4B    IN................TFNTVEYVMYMMLLLTAYFLNHL...................

           50         60        70        80        90        100
KpPno1    YTGHSAVSSTFN..ASSVVSDNKINGYGLPLIDTESNSRYEDPDDISIENELR.YRIAQS
KpMn4A    DLANSSLSISD.....FIKLKLK............GQKTYHKFDEHVFA.....AMARIQ
OpoMnn4   RATSSSTDPDD.....FIKSFEDQ..............................LDSLDI
ScMn4     .PNYNELDSDDDSWYSILTSSFK.............NDRKIQF.......AKTLYENL..
ScMnn14   .........DGNWKSFMALFFK......PLAYTNRNNNHASFDLRSKDNVAKLYEKM..
YlMpo1    ......LNLTDK.LPDGVKSRVA...............................YMTDVG
KpMn4C    ........SLD.....ILGSHVK...............................YLPLRE
OpoAbv1   ................FVMSKNK...............................Y.....
KpMn4B    ......LHSLDN.INHLVESDVN...............................YQLLQR

             110       120       130       140       150       160
KpPno1    TKEEENMWKLDTTLTEASLKIPNIQSFELQPFKERLDNSLYNSKNIGNFYFYDPRLTFSV
KpMn4A    SNENGKLADYESTSSKTDVTIQNVEL.....WKRLSEEE..........YTYEPRITLAV
OpoMnn4   SQNYGKYWKFSTTLNNAVVTFD......PSDYKTTRESR..........LLYDPRITMSV
ScMn4     KFGTNPKWVNEYTLQNDLLSVK......MGPRKGSKLESVDELK....FYDFDPRLTWSV
ScMnn14   NFDTSGKWIDTYTLKNNLLTVK......MGPEKGQVLDSVDELR....YYDNDPRLVWSV
YlMpo1    LVSGGA..RSKAMAGNSSVRISHVPLT....KIKFAENE..........KEFNPKWAQKK
KpMn4C    KVDPEEALH....LHGLDLSVAELPF.....FNDDMMSE..........FNYDPRLPTAL
OpoAbv1   EVDSEASFRKLQKSHTNDLAYV......ILPDEKHTEPHAPSK......SLFDQRVAVAA
KpMn4B    VTNKVKLFDEEAVL..........PF.....AKNLNRRT..........ERFDPRLPVAA

              170             180       190                  200
KpPno1    YLKYIKDKLASGSTT.......NLTIPFNWAHFRDLSSL...........NPYLDIKQ..
KpMn4A    YLSYIHQRTYDRYATSYA..PYNLRVPFSWADWIDLTAL...........NQYLDKTK..
OpoMnn4   YLSHIRRQLLEQGSGNAT..LAKIAVPFSWQDWVDLSVL...........NRYLELPE..
ScMn4     VLNHLQNNDADQPE..........KLPFSWYDWTTFHEL...........NKLISIDK..
ScMnn14   LLDHLLESDSN.............EYAFSWYDWANFDST...........NKLIALRH..
YlMpo1    ALDS..................SSDYSFEWKDWVDLTEV...........TGLFRAIEVG
KpMn4C    ILKLVLDHISVRNGTFDA....KFKVPFNWKLWVDLHSRLVPS.......NSWYNRFRLP
OpoAbv1   WLSNLKKHLMDNNGVLD....PDFQLPFSWAEWVDLESKLTLDKRHLHLWKSYHPEMQNR
KpMn4B    YLRSLQDQYSELPQGTDLNDIPPLEVSFHWDDWLSLGIA...........STFWDAFD..

               210                          220        230
KpPno1    .EDKVACDYFYESS....................NKDKRKPTGNCIEFKDVR....DEHL
KpMn4A    .....GCEAVFPR..........ESEATMKLNNITVVDWL.EGLCITDKSLQNSVNSTYA
OpoMnn4   .DQRPTCDDILYQ..........GPAYNPEHKFKNPNRNI.DRACVDNDRYL........
ScMn4     ..TVLPCNFLFQSAFDKESLEAIETELGEPLFLYERPKYA.QKLWYKAARNQDRIKDSKE
ScMnn14   ..TNISCQFVCEGAFDKNVLEMVESEVQEPLFVTNRNKYD.ESLWYNRVR...KVVDSNS
YlMpo1    .....SCG....................................................
KpMn4C    SGRFETCDEF......K........................RFFGITKNHFGTDLDNCVD
OpoAbv1   TFSDLTCADF................................RLLYADKDEV..LQVCSE
KpMn4B    ..................................NYNKRQ.GENAISYEQLQ........
                                                       *
```

```
         240                                250                         260            270                 280
KpPno1   IQY.........GISSKDHLPG......PFI.LKSLGIPMQHTAKRLESNLYLLTGAPVP
KpMnn4A  EEI...........NSRDILSP......NFHVFGYSDAKDNPQQKIFQSKSYINSKLPLP
OpoMnn4  ............GGTRRALLP.......GFN.FKDRTGKKSFRDKLIHAKSYLLSYAPVP
ScMnn4   LKK....HCSKLFTPDGHGSPKGLRFNTQFQ.IKELYDKVRPEVYQLQARNYILTTQSHP
ScMnn14  VQQAIHDHC.....MNNDAYSNGTPFELPFI.ISEISERLRPEVYDLQAKNHLLYSNFTP
YlMpo1   ..............KNKQCLS.......KLQVTGPPTQVEPQAFFNRVGKVFLDQTMPKP
KpMnn4C  IEY...........DTPEGYP.......KFKVLHAEDKALPYEARIIYGASYLYHEAQNP
OpoAbv1  LSH...........EERTQYP...WYPFSFKITDRTGSELAEAGRILFGASYMLTTAPPP
KpMnn4B  ..........................AIL.VNDLEDFSPYTAHILHSNVEVYKYRTIP

                  290            300
KpPno1   LSLSFM...TKKGLYQVGVDQTGK..................................
KpMnn4A  KSLIFL...TDGGSYALTVDRTQ...................................
OpoMnn4  TSIVFL...TENGSYEVNISGDV...................................
ScMnn4   LSISII..ESDNSTYQVPLQTEK...................................
ScMnn14  LSLTVL..DSDKDAYRINLKTTDS..................................
YlMpo1   KQLIYLTEKESRGKREEPVEIES..HVPVVREPELGDFSRSRDAKSIQNAKREATEEATG
KpMnn4C  KRLIFLGLGKSNESLILPVEAND...................................
OpoAbv1  ARVHFIGAGLNKSSVVLLTERDAVTHMPI.............................
KpMnn4B  QKIVYM...SNKGYFELLVTEKEK..................................

                     310                           320                           330
KpPno1   LDPN......IARTELWEFYK........NGKENL..........QFNAQEELSH..LIE
KpMnn4A  .NKR......ILKSGLLSHFF........SKKKKEHN..LPQDQKTFTFDPVYEFNRLKS
OpoMnn4  ...N......MLDSGLFDEFV........ELQANE..........TVVADASLEHEMLVN
ScMnn4   .SKN.....LVQSGLLQEYINDNINSTNKRKKNKQ.............DVEFNHNRLFQ
ScMnn14  SKSN......IVQTNLLQNYI........KRHRNE..........MVNGDLIFNHTSMFE
YlMpo1   ESANEGQSRDSARASARDIAF........SEADPVHEDSYDDDENGTILVPTAESEEELQ
KpMnn4C  .SSN......LMQFN..HEYA......................RSFNDQPFVSLEELVK
OpoAbv1  .SSNR..SRDIYSIG..AELVSNEARLRGVSVDRVHKDGVRIDQMVSELSTLFNYDTSVQ
KpMnn4B  ....  ......LSNEGLWSIFH........QKQGGLNE..............FSSLNLIE

            340                        350               360                   370            380
KpPno1   TVPSSSNSSSG........EGYFTTE..LK.ENNF....ELPLSKNDFTFDDSEVESLIK
KpMnn4A  QVKPRPISS..........EPSIDSA..LK.ENDY....KLKLKESSFIFNYGRILSNYE
OpoMnn4  SVSPSKS............ELDFDSL..FTAEKNY....EFVIPPDRFNYSYDQIIENYE
ScMnn4   EFVNNDQVNS.....LYKLEIEETDK..FTFDKDL.....VYLSPSDFKFDASKKIEELE
ScMnn14  KFLHHGSTKK......RKLDVEALDK..TIYAGEY.....LELSPSDFQFNAKERIIELE
YlMpo1   EYADKDAAA..........KSYLRSG..WSRGQKY.....VELPRELFTWDIHEEIDKGL
KpMnn4C  KVSLTLNLNSDKVLPINELDVIKDTPRLMNHNNQG......LSIDKSSFQWDLERELQLLE
OpoAbv1  TIYGED............ELVFDKSYFLKLDDQVKVMTSLEMTKQDFVYEHEEIMNAMK
KpMnn4B  EV..............DALDEIYDSKGLPAWDPPFP..EELDASDEDFKFNATEELAKVE

                   390                       400            410              420
KpPno1   ...GLSEQ.DLDLH.......TQRYKESLQYSFATREN..DVKKYFYEARMIINTVN...
KpMnn4A  ..ERLESLNDFEKS..........HYESLAYSSLLEAR..KLPKYFGEVILKNP......
OpoMnn4  ..KRIDEL...DEK.......QLRHLQTLKYSRSIPST..KLKKSFREVNINWPATYNGH
ScMnn4   EQKKLYPD.KFSAH.......NENYLNSLKNSVKTSPA..LQRKFFYEAGAVKQY.....
ScMnn14  ..TRLRSE.GLPSH.......DTHYLRSLKTSVNTSPA..LQQKYFAEASDITDA.....
YlMpo1   TKKSVDSD.DPSRE.......QVAHSQFLSQHWKHIK..KSGKHFSEAWVVGDT.....
KpMnn4C  ..HRTSQVNDVEGL.......DAGIYSTIQCEMRSMY...DFSKYFHESKVSGKY.....
OpoAbv1  ..QRIEKQLETKPDTVVDSLFDHLLLKNIEREMDLFEANGRHKPYLREAYVNN.......
KpMnn4B  ...QIKEP.KLE.........DIFYQEGLQHGIQTLPS..DASVYFPVNYVENDP.....
                                       *

              430            440                450                460            470            480
KpPno1   ..KEGGAHYDWRFFNGAMNHESSGFTEEERQLRKRSVLHRLLRNWLVFNYQQGSPTWLAH
KpMnn4A  ..QDGGIHYDYRFFSGLIDKTQINHFEDETE.RKKIIMHRLLRTWQYFTYHNNIINWISH
OpoMnn4  KVTENGGHYDFRFFNGFVTESKLNEYDDVNE.KRKIMLHRIIHTWLQFTYKEGIVSFLAH
ScMnn4   ..KGMGFHRDKRFFNV...DT...LINDKQE..YQARLNSMIRTFQKFTKANGIISWLSH
ScMnn14  ..TADGHHRDRRFFSI..GHN...LLNDPQE..FEARLNSLIRNFQKFVKANGLISWLSH
YlMpo1   ..KAAGVHYDWRFFSE.........LNTIDE..KRVILRKLVRAWLDFTSREGIITWLAH
KpMnn4C  ..LPSGEHYDWRFFNGFYL.......SQQE..NLAVLHRLGRAWLRFSRAAGLHTWIAH
OpoAbv1  ..THLGAHFDWRFFNG.........IDHPQE.YRKAIIHRIGRAWLRFCYQSGFRTFVAY
KpMnn4B  ..GLQSHHLHFPFFSGMVL.........PRE..IHSSVHHMNKAFFLFARQHGYVVWFFY
            *                          **
```

```
                 490       500       510       520       530            540
KpPno1    GTLLSWYWNSLMFPWDYDIDVQMPIKSLNNLCANFNQSLIIEDLTE.....GYSSFFLDC
KpMnn4A   GSLLSWYWDGLSFPWDNDIDVQMPIMELNNFCKQFNNSLVVEDVSQ.....GFGRYYVDC
OpoMnn4   GTLLSWYWNALVFEWDNDIDVQMPIMDFDRFCMKYNNSLIVEDVQH.....GYGKYYVDC
ScMnn4    GTLYGYLYNGMAFPWDNDFDLQMPIKHLQLLSQYFNQSLILEDPRQ.....GNGRYFLDV
ScMnn14   GTLYGYLYDGLKFPWDVDHDLQMPIKHLHYLSQYFNQSLILEDPRE.....GNGRFLLDV
YlMpo1    GTLLGWYWNGQSLPWDFDGDVQMPIREFDRFARLYNQSLVIDES.......AGGRYYVDV
KpMnn4C   GTLLGWYWNGLILPWDQDLDVQMTVQSLYLLGRNFNSSLVTDVSIEDGYSSALGHYYIDV
OpoAbv1   GSMLGWIRNGLTLPWDEDIDVVVSVDSLYKIARNHNQTLIVDVSSEDKYAAGIGSYLLDI
KpMnn4B   GNLIGWYYNGNNHPWDSDIDAIMPMAEMARMAHHHNNTLIIENPHD.....GYGTYLLTI
             *                           *                          *

                     550       560       570       580       590
KpPno1    GSSITHRTKGKGLNFIDARFINVETGLYIDITGLSTSQSARPPRFSNASKKDP.......
KpMnn4A   TSFLAQRTRGNGNNNIDARFIDVSSGLFIDITGLALTGSTMPKRYSNKLIKQPK......
OpoMnn4   GPYPTHRTKGNGRNNIDARFIDVDSGMYIDITGLALTDTIKIPPRLERLDRQRKANN...
ScMnn4    SDSLTVRINGNGKNNIDARFIDVDTGLYIDITGLASTSAPSRDYLNSYIEERLQEEHLDI
ScMnn14   GSAITVGVHGNGENNIDARFIDIDSGIYIDITGLSVSSDAAKQYMSKFVEEESSGESFSA
YlMpo1    GPSYVERLRGNGKNVIDARFIDVDSGMYIDITALAYAEQ.....................
KpMnn4C   GSSFFVRDKLNGNNAIDARFVDTETGLYVDITALAFTDHLKLKLTTKEKVELQK......
OpoAbv1   GPSFYSRVRGVGHNAIDGRMIDTMSGVYVDITAIAWTPDYFSQHNIDDVVR.........
KpMnn4B   SPWFTKKTRGG..NHIDGRFVDVKRGTYIDLSAISAMHGIYPDWVRDGVKENPK......

KpPno1    ............................................................
KpMnn4A   ...........KSTDS............................................
OpoMnn4   ........EQGKSEDA...........LPAEQTEGLSDP.......GASRNVKRAPVKSN
ScMnn4    NNIPESNGETATLPDKVDDGLVNMATLNITELRDYITS........DENKNHKRVPTDTD
ScMnn14   .LIEDYKFDENDYFDEV.DGREGLAKYTIHELMEWVNSHPD.....DFTDAEKNLVTKT.
YlMpo1    ............................................................
KpMnn4C   ............VMDP...........NVKEKLQWIKNKYSTATLPGVIETDRNKVSDA.
OpoAbv1   ...........TLVDP...........EYPSKIDKVKDK..................DS.
KpMnn4B   ............................................................

                                                   600       610
KpPno1    ..................................IYNCRNNHFYSHNNIAPLKYTLMEG
KpMnn4A   .............TGSTPENGLTRNLR..QNLNAQVYNCRNGHFYQYSELSPLKLSIVEG
OpoMnn4   ....KGPERS..PEALE............RNKQLQIYNCRNNHSYTYSELSPLKLSMMEG
ScMnn4    LKDLLKKELEELPKSKTIENKLNPKQRYFLNEKLKLYNCRNNHFNSFEELSPLINTVFHG
ScMnn14   ....YKKELA.ISRSDYAEKDLSPKQRYLVNEKYNLYNCRNQHFSSLNIISPLRNTMFSG
YlMpo1    ................................QEKFHCKNWHRYELESVSPLRRTLFEG
KpMnn4C   ....LEKQFH..............DFKFDNFVNKELFHCRNNHFYKYGEVGRLRSTMFEG
OpoAbv1   ....YREQID..NKARD............MQRNHEIYHCRNDNVFRLDELTPMVPTYFEG
KpMnn4B   ................................NLALADKNGNWYLTRDILPLRRTIFEG
                                                                 *

          620       630       640       650       660
KpPno1    VPSFIPQQYEEILREEYT.TGLTSK....HYNGNFFMTQLNLWLERDPML..........
KpMnn4A   ALTLIPNDFVTILETEYQRRGLEKN....TYAKYLYVPELRLWMSYNDIY..........
OpoMnn4   SPCLIPHDFATVLNSEYN.GGLRKK....HFNNHLFIDELRFWVSTATLQQHAYALSGHL
ScMnn4    VPALIPHRHTYCLHNEYH.VPDRYAF..DAYKNTAYLPEFRFWFDYDGLKKCSNINSWYP
ScMnn14   VSAFVPNRPIATLNNEYK.VPAKYGL..LSFQGKVYLPEFRYWFSFADMKKFANLQ....
YlMpo1    KEAYIPNNFESILNQEYKKAPLVNT....RFEGHFWNKFIKMWVQQDQCE..........
KpMnn4C   VPALIPFEFESILKREYP.KGLTLK....HFSNHFWDPVNRLWVPEKKKK..........
OpoAbv1   VRTHVPHKFKQILDRKYP.RALQRITEPENQPFRTYKKNLRLWVDDAQCP..........
KpMnn4B   SRSYTVKDIEDTLLRNYGDKVLINT....ELADHEWHDDWKMWVQKKKYCTYE.......

                    670       680                  690
KpPno1    ...........ALVPSSKYEIEGGGVDHNKI.......IKS......ILELSNIKKLELL
KpMnn4A   ...................DILQGTNSHGRP.......LSAKTMATIFPRLNSDINLKKF
OpoMnn4   ...............VKNVDLRKFYNSHKLP......................MLLKM
ScMnn4    NIPSINSWNPNLLKEISSTKFESKLFDSNKVSEYSFKNLSMDD.......VRLIYKNIPK
ScMnn14   ............LKEPKITRLESPLND...........LKFSD.......ISLLITNILK
YlMpo1    ...................MLQIEENVDQRA.......VRENGEP..........TTFGA
KpMnn4C   ......................................IRH......I............
OpoAbv1   ........................GNDNDGL............................L
KpMnn4B   ...................EFEDYLSAHGGVE.....YDEDGVL..........TLEGA
```

```
           700       710       720       730       740       750
KpPno1     DDNPDILEEVIRTYELTSIHHKEMQYLSSVKPDGDRSMQSNDITSSYQEFLASL.KKFQP
KpMnn4A    LRNDHTFKNIYSTFNVTRVHEEELKHLIVNYDQNKRK......SAEYRQFLENL.RFMNP
OpoMnn4    LGTDELLKEFYLTHKTTKMHLEEFSLM..NQLSDPQK......KERYYSIMEEYKRTHDP
ScMnn4     AGFIEVFTNLYNSFNVTAYRQKELEIQYCQNLTFIEK......KKLLHQLRINVAPKLSS
ScMnn14    CGFHSVFASLFNSFDSTVYRLKELEIQYDPSLSEEEK......SSLLKTLRRGMSKKIKS
YlMpo1     CYRPEYLKRYHETHKMSKAHEGEMEAIRQK.............ADVWEWIREEF......
KpMnn4C    ........EFSLTKEVTESHKKELAQIHGNET.........GITSDF.........AYSP
OpoAbv1    CRDEDVLEEYAHTKEYTARHLELLKHIYDNALSIEDE....TVPMRYDQFIVRYAQLLNA
KpMnn4B    CGFEEVRQDWIITRESVNLHMKEWEAIQRNES...........TTEYTA......KDLPR

             760
KpPno1     LRKDLFQF....................................................
KpMnn4A    IRKDLVTYESRLKALDGYNEV.......................................
OpoMnn4    MRKSLHGYISEVVALGGYHPQ.......................................
ScMnn4     PAKDPFLFGYEKAMWKDLSKSMNQTTLDQVTKIVHEEYVGKIIDLSESLKYRNFSLFNIT
ScMnn14    PEKDPIIYIYERKLWENVEKLLNASNIYNIASQVEKEKGKEFVERSQQVYERNFDGFRLP
YlMpo1     ............................................................
KpMnn4C    FRIDPWLSRYRKKMT.............................................
OpoAbv1    ............................................................
KpMnn4B    YRPDSF.....KNLLDGVSNHGNG....NVGKI...........................

KpPno1     ............................................................
KpMnn4A    ............................................................
OpoMnn4    ............................................................
ScMnn4     FDETGTTLDDNTEDYTPANTVEVNPVDFKSNLNFSSNSFLDLNSYGLDLFAPTLSDVNRK
ScMnn14    DGGNSKTVN...............................DLNSKGLNLFG....DNKKT
YlMpo1     ............................................................
KpMnn4C    ............................................................
OpoAbv1    ............................................................
KpMnn4B    ............................................................

                                   770
KpPno1     ......................ERIDLSKH..............................
KpMnn4A    ......................EELEKKQENREKERKEKKEKEEEKEKKEKEEEKEKKEKE
OpoMnn4    ......................EEPARSALSETSEELDRVVITGKSQSNAGARRIQK..
ScMnn4     GIQMFDKDPIIVYEDYAYAKLLEERKRREKKKKEEEEKKKKEEEEKKKKEEEEKKKKEEE
ScMnn14    SNNIFGSD....................................................
YlMpo1     ............................................................
KpMnn4C    ............................................................
OpoAbv1    ............................................................
KpMnn4B    ......................EHVKLE................................

KpPno1     ............................................................
KpMnn4A    EKEKKEKEEKERKEKEEKEEYEEDDNEGEQPTEQKSQQE.....................
OpoMnn4    ............................................................
ScMnn4     EKKKKEEEEKKKKEEEEKKKQEEEEKKKKEEEEKKKQEEGEKMKNEDEENKKNEDEEKKK
ScMnn14    ............................................................
YlMpo1     ............................................................
KpMnn4C    ............................................................
OpoAbv1    ............................................................
KpMnn4B    ............................................................

KpPno1     ...........................................RKQ
KpMnn4A    ...........................................AKE
OpoMnn4    ...........................................SPT
ScMnn4     NEEEEKKKQEEKNKKNEDEEKKKQEEEEKKKNEEEEKKKQEEGHSN
ScMnn14    ...........................................QKY
YlMpo1     .............................................E
KpMnn4C    ...........................................RSQ
OpoAbv1    ...........................................RLE
KpMnn4B    ...........................................HND
```

Figure S4. Multiple alignment of amino acid sequences of Mnn4 homologs from *O. polymorpha* (Opo), *K. phaffii* (Kp), *S. cerevisiae* (Sc), *Y. lipolytica* (Yl), generated using MAFFT and visualized in ESPript with BLOSUM62 matrix. Red background indicates strictly conserved residues; blue boxes highlight position with physicochemical similarity. Asterisks mark positions distinguishing Mnn4 from Abv1 protein groups.

Table S1. Oligonucleotides

| Name | 5'-3' sequence |
|---|---|
| ABV2U | CGAGTTGGAAGGAGAAGAC |
| ABV2L | GCTGCCTGGTTTGGATTG |
| OpoMNN4AU1 | CACATCAAACCCGCTCGCTG |
| OpoMNN4L1 | CCCGTTTGCCACTGTGTGCTG |
| OpoMNN4AL1 | GTGCTCTACAATCTCCTGTCC |
| OpoMNN4U1 | CTCACAAGACCACCAAGATGC |
| OpoOCH1U1 | GAAGACAAGATCGAGTGGGA |
| OpoOCH1L1 | GTTCTTCTGGATCTCGCTTG |
| OpaOCR1AL | CGGTGGTGAACGCGGTGGTG |
| OpaOCR1AU | GGGGCACAGATGACAAAATG |
